# β-arrestin1 directly engages Gαs to sustain endosomal GPCR signaling

**DOI:** 10.64898/2026.09.27.754863

**Authors:** Qian He, Qingning Yuan, Xinheng He, Wen Hu, H. Eric Xu, Li-Hua Zhao

## Abstract

β-arrestins (βarrs) are canonical terminators of G protein–coupled receptor (GPCR) signaling, yet internalized receptors paradoxically sustain βarr-dependent G_s_/cAMP signaling from endosomes. How a desensitizing scaffold can instead promote G protein activation has remained unresolved, because the receptor–βarr–G protein assemblies described so far position the two transducers on opposite faces of the receptor, with no contact between them. Using the parathyroid hormone type 1 receptor (PTH1R), we show that βarr1 binds Gα_s_ directly, engages its nucleotide-free state, and does so far more strongly than it engages Gβγ. A cryo-EM structure of the agonist-bound PTH1R– βarr1–Gα_s_ complex at ∼3.1 Å resolution reveals how: βarr1 anchors to the phosphorylated receptor C-tail, leaving the transmembrane core free for Gα_s_, and reorients to build a direct, three-node interface in which its finger, C- and back loops clamp the Switch I, Switch II and α3/β5 regions of Gα_s_. Disrupting either side of this interface abolishes complex assembly and selectively collapses the internalization-dependent phase of cAMP production, and the same interface governs βarr1–Gα_s_ coupling at the glucagon-like peptide-1 receptor. β-arrestin1 therefore acts as a positive regulator that holds Gα_s_ in a signaling-competent state, providing a unifying structural basis for sustained endosomal GPCR signaling.

## Introduction

Signal transduction by G protein–coupled receptors (GPCRs) has long been described in binary terms. Agonist-activated receptors at the plasma membrane engage heterotrimeric G proteins to initiate intracellular signaling, after which GPCR kinases (GRKs) and β-arrestins (βarrs) terminate that signaling by sterically occluding the G protein-binding cavity and driving receptor internalization^1–6^. In this view, G proteins and βarrs are mutually exclusive transducers, and internalization marks the end of receptor signaling.

That view was overturned by the discovery that many internalized receptors continue to generate G protein-dependent cAMP from endosomes, producing a second wave of signaling that is spatially and temporally distinct from the plasma-membrane response^7–11^. Critically, this sustained endosomal signaling is predominantly βarr-dependent ^12^—with βarr, the very scaffold long thought to terminate it — and drives physiological and pharmacological outcomes that the classical desensitization model cannot explain. Because endosomal signaling underlies the prolonged efficacy of clinically important receptor agonists, how it is sustained carries direct therapeutic significance.

The parathyroid hormone type 1 receptor (PTH1R), a class B GPCR essential for calcium and phosphate homeostasis, is the prototypical example of this noncanonical behavior^8,13–16^. Upon stimulation, PTH1R first activates G_s_ at the plasma membrane to produce a transient cAMP pulse; following βarr-mediated internalization, it continues to signal through G_s_ from endosomes, generating a second, sustained phase of cAMP that governs its pharmacological actions^13,17^. This spatiotemporal pattern, now validated across diverse GPCRs^10,11^, poses a fundamental paradox: how can an internalized receptor sustain G_s_ signaling while simultaneously engaging βarr, a transducer classically associated with receptor desensitization?

Structural studies have made such coexistence conceivable. βarr engages receptors in two distinct modes: a core conformation, in which βarr inserts into the transmembrane cavity and directly competes with G protein binding^18–27^; and a tail conformation, exemplified by the glucagon receptor (GCGR)-βarr1 complex^28^, in which βarr anchors to the phosphorylated receptor C-tail, leaving the transmembrane core free for G protein coupling. The tail-engaged mode therefore raises the possibility that a receptor, βarr, and G protein could occupy a single assembly to sustain endosomal signaling.

Consistent with this idea, several studies have proposed receptor–βarr–G protein assemblies that support prolonged endosomal signaling^9,29–32^, and cryo-EM has established their structural feasibility, capturing a tail-engaged βarr1 in a β_2_-adrenergic receptor/vasopressin 2 receptor (β2V2R) chimera and a distinct architecture in an atazanavir-stabilized β2AR megaplex^9,30,31^. In these assemblies, however, βarr lies far from the G protein and makes no direct contact. The central question therefore remains unanswered: does βarr actively sustain G protein activation, or does it merely permit the two transducers to occupy the receptor at once? Whether Gα_s_ itself directly participates, how such complexes are organized, and how βarr contributes to sustained G protein signaling have remained unknown.

These questions are difficult to address because the relevant complex is transient, conformationally heterogeneous, and destabilized by the very nucleotide exchange it exists to enable. Here, combining transducer-complementation assays, cryo-EM, and structure-guided mutagenesis, we show that PTH1R, βarr1, and nucleotide-free Gα_s_ assemble into a cooperative ternary complex. Our ∼3.1 Å structure reveals a previously unrecognized, direct interface between conserved loops of βarr1 and Gα_s_ that is required both for complex stability and for the internalization-dependent phase of cAMP signaling, and that is shared across class B GPCRs. Together, these findings resolve a long-standing paradox by demonstrating that βarr1 directly promotes, rather than terminates, G protein activation after receptor internalization, and they establish a conserved structural mechanism for endosomal signaling.

## Results

### βarr1 directly engages nucleotide-free Gα_s_ downstream of PTH1R

Sustained PTH1R signaling requires βarr-dependent receptor internalization and delivery to endosomes, a process that depends on dynamin-mediated endocytosis. We first examined whether PTH1R can simultaneously engage both G protein and βarr1 to support endosomal signaling. Dyngo-4a, a dynamin inhibitor, effectively blocks receptor endocytosis without altering basal cAMP levels, enabling specific assessment of endosome-derived cAMP signaling^33^. In GloSensor cAMP accumulation assays, Dyngo-4a treatment attenuated cAMP production induced by LA-PTH (**Extended Data Fig. 1a**), a long-acting analog of PTH^34^, while having no significant effect on cAMP detection induced by forskolin (**Extended Data Fig. 1b**), which directly stimulates adenylate cyclase. Consistently, overexpression of a dominant-negative Dynamin2 mutant (K44E, Dyn2_K44E), which prevents vesicle scission^35,36^, also suppressed cAMP production compared with cells expressing wild-type Dynamin2, the ubiquitously expressed isoform that predominates in non-neuronal cells such as HEK293^35,37–39^ (**Extended Data Fig. 1c, left panel**). By contrast, Dyn2_K44E had little effect on cAMP accumulation mediated by the phosphorylation-deficient PTH1R-T/S to A mutant (**Extended Data Fig. 1c, right panel**), indicating that receptor C-tail phosphorylation is required to generate the dynamin- and βarr1-dependent, internalization-driven phase of sustained signaling. Manipulating βarr1 levels confirmed its central role: overexpression enhanced the sustained cAMP phase, and this enhancement was attenuated by co-transfection of a βarr1-specific siRNA (**Extended Data Fig. 1d**), consistent with a βarr1-dose-dependent contribution to sustained signaling. βarr1-dependent internalization is therefore a prerequisite for sustained PTH1R signaling.

If βarr1 supports rather than terminates G protein signaling in this compartment, it should engage the G protein directly. We tested this using NanoBiT complementation assays in which Large Bit (LgBiT) was fused to the N- or C-terminus of Gα_s_, Gβ, or Gγ, and Small Bit (SmBiT) to the N-terminus of βarr1 (**Fig. 1a**). Following PTH1R activation, robust complementation was detected between βarr1 and either miniGα_s_ (**Fig. 1b**)— a nucleotide-free Gα_s_ mimetic carrying engineered truncations and stabilizing mutations^40^— or Gβ (**Fig. 1c, d**; the latter showing a zoomed-in x-axis), but only when LgBiT was placed at their N-termini, and not when fused to their C-termini or to Gγ (**Fig. 1e**–**g**; **Extended Data Table 1**). This strict orientation dependence indicates a defined binding geometry rather than nonspecific proximity. The βarr1–Gα_s_ signal was substantially larger than βarr1–Gβγ coupling (**Fig. 1c, d**), and co-expression of Gβγ diminished it (**Fig. 1b**), identifying Gα_s_, rather than Gβγ, as the dominant βarr1-interacting G protein subunit.

**Fig. 1.**
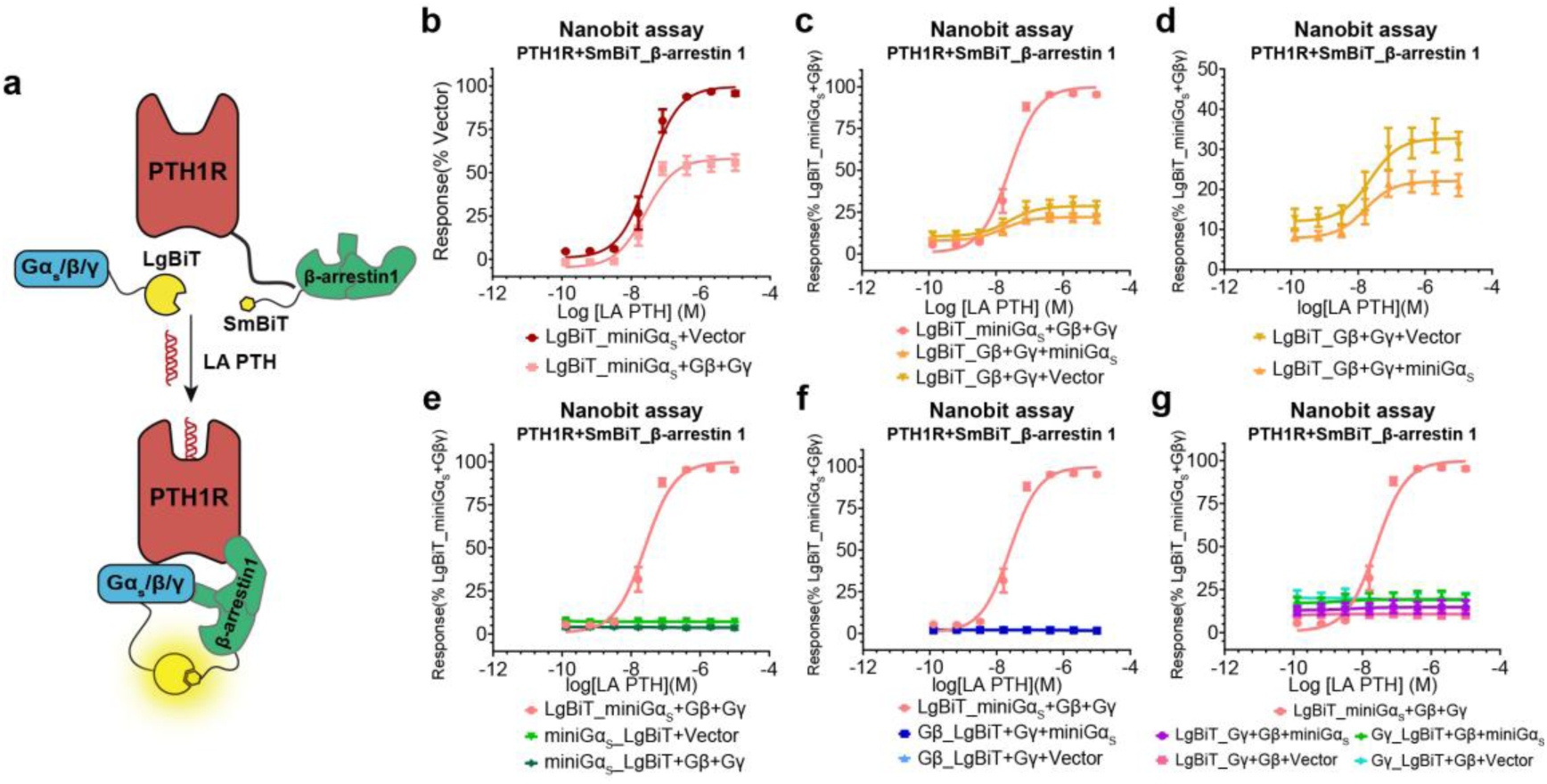
β-Arrestin1 directly engages a nucleotide-free state of Gα_s_ downstream of PTH1R. **(a)** Schematic of the NanoBiT complementation assay used to assess interactions between βarr1 and G protein subunits downstream of PTH1R activation. Unless otherwise indicated, all assays in **(b)**–**(g)** were performed in cells co-expressing PTH1R and SmBiT–βarr1, with NanoBiT responses measured following LA-PTH stimulation; ‘Vector’ denotes replacement of the indicated co-expressed G protein subunit (Gβγ or miniGα_s_) with empty vector. **(b)** NanoBiT dose**-**response curves of βarr1 interaction with miniGα_s_, with LgBiT fused to the N-terminus of miniGα_s_, in the presence or absence of co-expressed Gβγ following LA-PTH stimulation. Luminescence signals were normalized to LgBiT–miniGα_s_ + SmBiT–βarr1 + Vector controls. **(c)** NanoBiT concentration-response curves measuring the interaction between βarr1 and Gβ, with LgBiT fused to the N-terminus of Gβ, in the presence or absence of co-expressed miniGα_s_ following LA-PTH stimulation. Luminescence signals were normalized to the LgBiT–miniGα_s_ + SmBiT–βarr1 + Gβγ control. **(d)** Comparison of the βarr1–Gβγ interaction in the presence or absence of miniGα_s_, shown with a zoomed-in x-axis corresponding to the data in Fig. 1c. **(e)** NanoBiT complementation assays measuring the interaction between βarr1 and miniGα_s_, with LgBiT fused to the C-terminus of miniGα_s_, in the presence or absence of co-expressed Gβγ. Luminescence signals were normalized to the LgBiT–miniGα_s_ + SmBiT–βarr1 + Gβγ control. **(f)** NanoBiT complementation assays measuring the interaction between βarr1 and Gβ, with LgBiT fused to the C-terminus of Gβ, in the presence or absence of co-expressed miniGα_s_. Luminescence signals were normalized to the LgBiT–miniGα_s_ + SmBiT– βarr1 + Gβγ control. **(g)** NanoBiT complementation assays measuring the interaction between βarr1 and Gγ, with LgBiT fused to either the N- or C-terminus of Gγ, in the presence or absence of co-expressed miniGα_s_. As neither orientation yielded detectable signal, both are shown together. Luminescence signals were normalized to the LgBiT–miniGα_s_ + SmBiT– βarr1 + Gβγ control. All the data are presented as mean ± S.E.M. from at least three independent experiments, each performed in triplicate.

We next asked which nucleotide state of Gα_s_ βarr1 recognizes. βarr1 bound robustly to two structurally independent nucleotide-free forms of Gα_s_: the minimized mimetic miniGα_s_ and the full-length dominant-negative mutant G112 ^40–42^ (**Extended Data Fig. 1e**). That βarr1 recognizes both a truncated and a full-length nucleotide-free Gα_s_ indicates that this interaction reflects the nucleotide-free state itself rather than any single construct. Assembly further required a phosphorylated receptor: substituting all Ser/Thr residues in the PTH1R C-terminal tail with Ala markedly reduced both βarr1**–** miniGα_s_ association and sustained cAMP production (**Extended Data Fig. 1f**), identifying receptor C-tail phosphorylation as a prerequisite for assembling the complex.

Together, these data define a PTH1R**–**βarr1**–**Gα_s_ complex in which the phosphorylated receptor licenses βarr1 to engage nucleotide-free Gα_s_, sustaining G protein signaling after receptor internalization.

### Cryo-EM structure of the PTH1R–βarr1–Gα_s_ complex

To resolve how βarr1 and Gα_s_ engage PTH1R simultaneously, we determined the cryo-EM structure of the PTH1R**–**βarr1**–**Gα_s_ complex. Because the assembly is transient, we combined three stabilization strategies: the high-affinity vasopressin type 2 receptor C-tail (V2RT) was appended to enhance βarr1 binding^9,30,43^; βarr1 was fused to the receptor through a flexible GSA linker; and the active βarr1 conformation was locked with the conformation-specific antibody fragment scFv30^4,43,44^. This chimeric construct retained wild-type-like behavior in both NanoBiT and cAMP assays (**Extended Data Fig. 1g**), and was co-expressed with miniGα_s_ and GRK2 to ensure receptor phosphorylation and efficient assembly.

From 111,056 movies we obtained a final reconstruction at a global resolution of ∼3.1 Å (**Fig. 2**, **Extended Data Fig. 2–4**, and **Extended Data Table 2**). The density map shows both transducers engaged at once: LA-PTH occupies the orthosteric pocket of PTH1R, while βarr1 adopts a tail conformation anchored to the phosphorylated receptor C-terminus (**Fig. 2**). Critically, this arrangement leaves the transmembrane core fully accessible, allowing miniGα_s_ to insert its α5-helix in the canonical activation mode (**Fig. 2b**). Focused refinement of PTH1R, miniGα_s_, βarr1, and scFv30 yielded a composite map for detailed interface analysis (**Extended Data Fig. 2e, 3**). The αN-helix of miniGα_s_ was not resolved, consistent with its dispensability for complex formation (**Extended Data Fig. 1h**) and with its known flexibility in the absence of Gβγ, as previously observed in the isolated Gα_s_ structure (PDB: 1AZT)^45^.

**Fig. 2.**
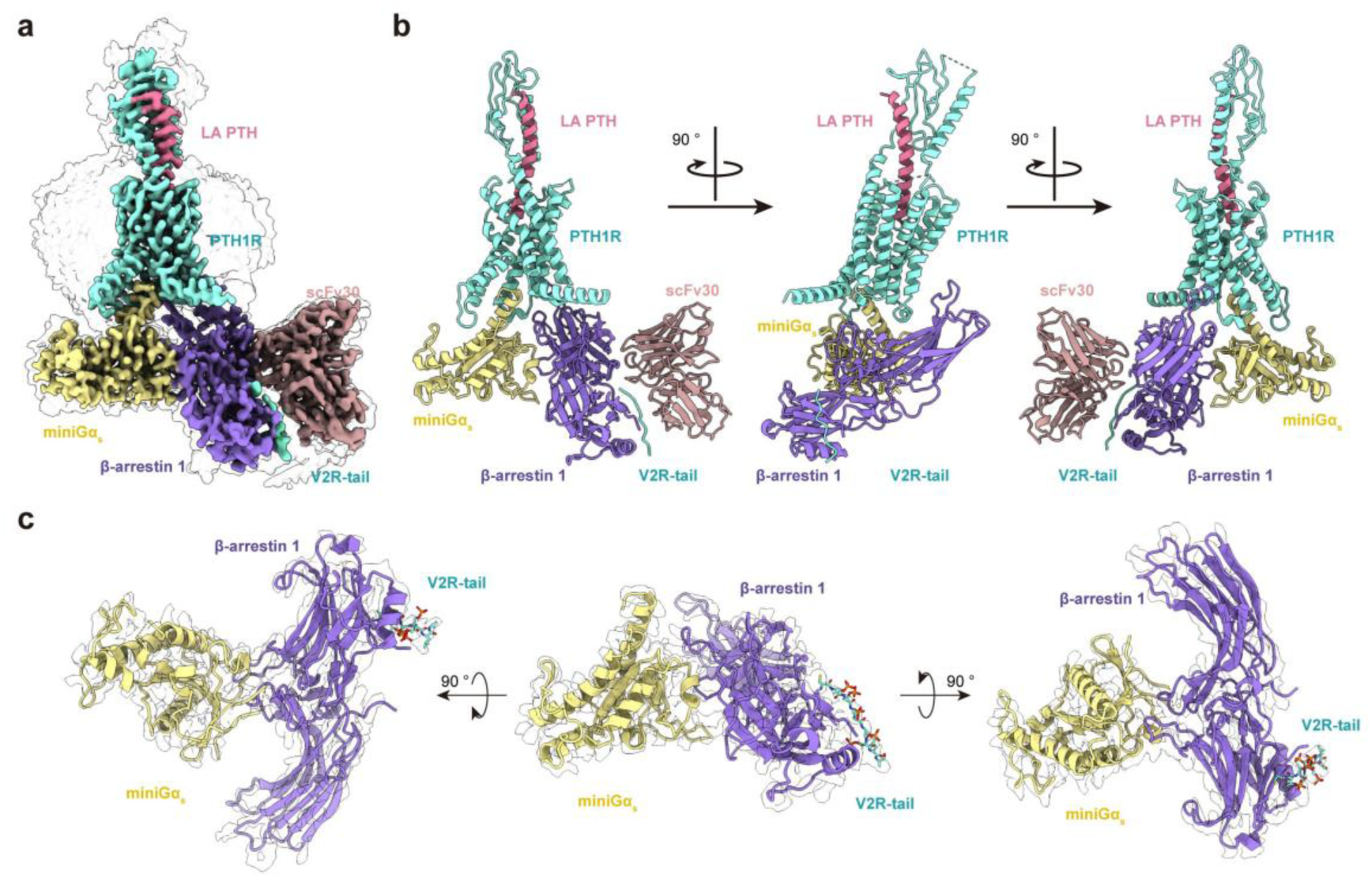
Cryo-EM structure of PTH1R–β-arrestin 1–Gα_s_ complex. **(a**, **b)** Cryo-EM map **(a)** and model **(b)** of LA-PTH–bound PTH1R–βarr1–miniGα_s_ complex shown in multiple orientations. LA-PTH, *Pale Violet Red*; PTH1R_V2RT, *Aquamarine*; miniGα_s_, *Kihaki*; βarr1, *Medium Purple*; scFv30, *Rosy Brown*. **(c)** Focused cryo-EM density map and corresponding model of βarr1 and miniGα_s_, displayed from various angles.

Hydrophobic clusters in the distal βarr1 C-lobe lie close to the lipid bilayer (**Fig. 2b**). To test whether membrane insertion drives assembly, we substituted residues within the βarr1 C-edge 197- and 344-loops^46^ with Gly or Ser. The 197-loop substitutions had minimal effect and the 344-loop substitutions only modest effects on complex formation (**Extended Data Fig. 1i**). Membrane anchoring may therefore stabilize the assembly, but it is not essential for it, pointing instead to the direct βarr1–Gα_s_ contact as the principal determinant of complex stability.

### Reciprocal structural adaptation accommodates both transducers

Accommodating both transducers at once requires coordinated structural adaptation across the assembly. The receptor maintains a canonical active conformation, with an outward displacement of TM6 (∼17.5 Å) relative to the tail-engaged GCGR–βarr1 complex^28^ (**Extended Data Fig. 5a–e**) that opens the intracellular cavity for miniGα_s_. The overall conformation of miniGα_s_ remains highly conserved compared with the binary PTH1R-Gα_s_ complex^47^ (RMSD 0.70 Å over 246 Cα atoms), but its interfacial regions undergo distinct localized shifts (**Extended Data Fig. 5f, g**); in particular, the intrinsically dynamic Switch II region^48^ is displaced by ∼5.2–8.2 Å to support inter-transducer coupling (**Extended Data Fig. 5f, g**).

The largest rearrangement occurs in βarr1, which adopts an orientation distinct from all previously reported GPCR–βarr1 complexes^22–24,28^ (**Fig. 3a, b**). Relative to the tail-engaged GCGR–βarr1 structure, βarr1 in the PTH1R–βarr1–Gα_s_ complex shifts ∼19.5° away from the receptor core and rotates ∼60.8° toward miniGα_s_ (**Fig. 3b**). This reorientation redirects the finger loop, back loop, and C-loop to present a complementary surface for direct miniGα_s_ engagement (**Fig. 3c–g**). Rather than competing for receptor occupancy, βarr1 and Gα_s_ therefore adapt structurally to bind cooperatively and simultaneously.

**Fig. 3.**
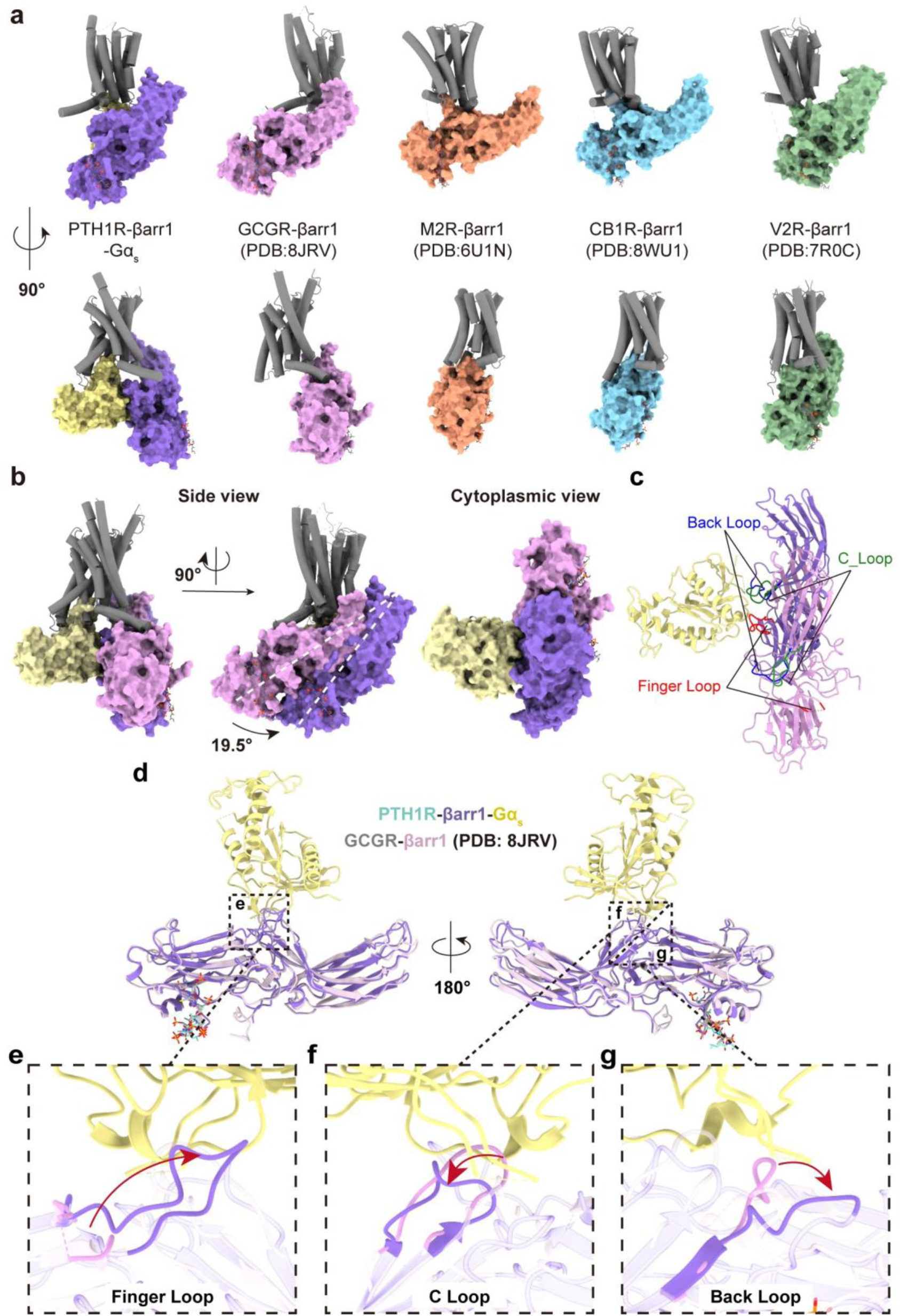
Conformational mode of β-arrestin1 in the PTH1R ternary assembly. **(a)** Structural comparison of βarr1 in the PTH1R–βarr1–Gα_s_ complex with previously resolved GPCR–βarr1 complexes, including GCGR–βarr1 (PDB: 8JRV), M2R–βarr1 (PDB: 6U1N), CB1R–βarr1 (PDB: 8WU1), and V2R–βarr1 (PDB: 7R0C). **(b)** Tail-engaged conformation comparison of βarr1 in the PTH1R–βarr1–Gα_s_ complex and GCGR–βarr1 (PDB: 8JRV), aligned by receptors. **(c)** Structural shifts of the finger loop, C-loop, and back loop in βarr1 compared between the ternary assembly and GCGR–βarr1 (PDB: 8JRV) , aligned by receptors. **(d)** Overall structural comparison of βarr1 from the ternary assembly and GCGR–βarr1 (PDB: 8JRV), aligned by βarr1. **(e-g)** Detailed conformational movements in the finger loop **(e)**, C-loop (**f**), and back loop (**g**) of βarr1 between ternary assembly and GCGR–βarr1 (PDB: 8JRV), aligned by βarr1.

This architecture differs fundamentally from previously resolved GPCR megaplexes, in which the two transducers remain spatially separated. In the β2V2R**–**βarr1**–**G_s_ megaplex^30^, βarr1 is positioned ∼77.6° further from the receptor core, precluding any G protein contact (**Extended Data Fig. 6a, b**). In the atazanavir-stabilized β2AR**–** βarr1**–**G_s_ megaplex^31^, βarr1 is repositioned to the opposite side of the receptor transmembrane domain (**Extended Data Fig. 7a–c**), with a ∼19.8° axial rotation and ∼31.8 Å displacement (**Extended Data Fig. 7d**). In both cases, local remodeling of the βarr1 finger, back, and C loops would be required to establish the direct βarr1**–**Gα_s_ interface seen here (**Extended Data Fig. 6c, d, and 7e, f**). Notably, Gα_s_ occupies a conserved position relative to the receptor across all three complexes (**Extended Data Fig. 6e, 7g**), with distinct electrostatic anchoring on the phosphorylated receptor tails accounting for the configurational plasticity of βarr1 (**Extended Data Fig. 6f**). Ternary assembly thus depends on mutual accommodation: βarr1 reorients and remodels its loops to receive Gα_s_, while Gα_s_ makes the localized interfacial adjustments that stabilize the interaction.

### A direct βarr1–Gα_s_ interface stabilizes the ternary complex

The central architectural feature of this cooperative assembly is a direct βarr1**–**Gα_s_ interface built from three interaction nodes, which are observed in our resolved structure (**Fig. 4a**) and further supported by an MD model generated from molecular dynamics simulations based on our structure (**Fig. 4b**). The first involves the βarr1 finger loop (residues 63**–**77), which comes into close apposition with a Gα_s_ surface spanning K233**–**W281 (**Fig. 4c**). The second centers on the βarr1 C-loop (residues 240– 249), which packs against a Gα_s_ surface spanning F208–V241 (**Fig. 4d**). The third involves the βarr1 back loop (residues 311–319), which is positioned adjacent to the Gα_s_ N66/S205 region (**Fig. 4e**). Because local density at this interface did not support unambiguous side-chain assignment, these loops were modeled at the Cα level, and the interface is therefore described here in terms of loop-to-surface proximity rather than discrete side-chain pairings. Together, the three nodes map onto the Switch I, Switch II, and α3/β5 regions that reorganize when Gα_s_ releases nucleotide.

**Fig. 4.**
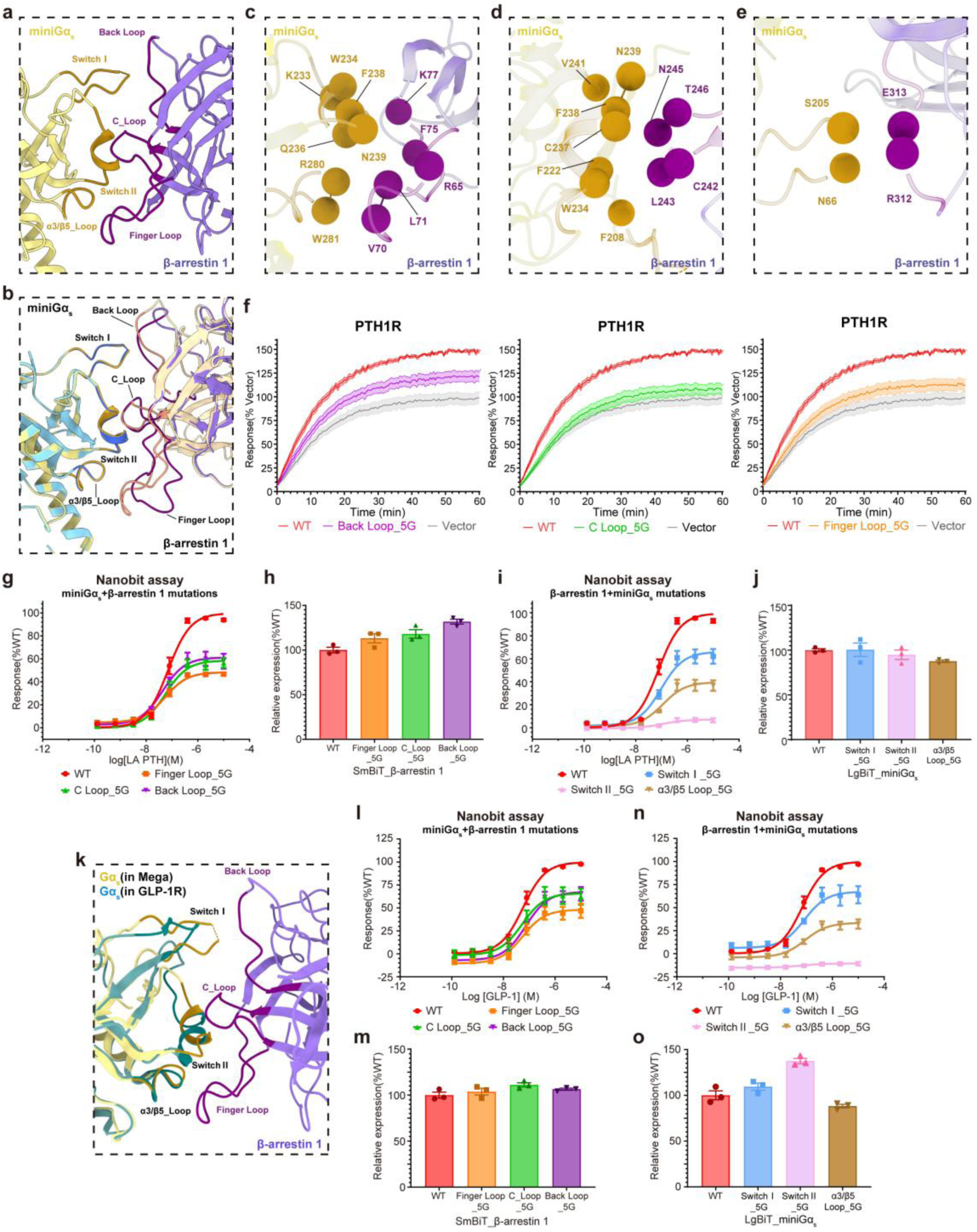
Structural basis and conservation of the β-arrestin1–Gα_s_ interface. **(a, b)** Structural overview of the interaction interface between βarr1 and miniGα_s_ in the complex, based on the cryo-EM model (**a**) and the molecular dynamics–refined model (**b**). **(c‒e)** Close-up views of the βarr1–miniGα_s_ interface highlighting contacts mediated by the βarr1 finger loop (**c**), C-loop (**d**), and back loop (**e**). Owing to limited local density at the interface, side-chain positions could not be reliably modeled; therefore, Cα atoms are shown as spheres to illustrate loop positioning and spatial organization. **(f)** Effect of substituting individual βarr1 interface loops with glycine linkers (5× Gly) on LA-PTH-induced sustained cAMP signaling in HEK293 cells expressing wild-type PTH1R. Luminescence signals were normalized to the response of vector-transfected controls. **(g**, **i)** NanoBiT complementation assays quantifying interactions between miniGα_s_ and wild-type or loop-substituted βarr1 (**g**) and between βarr1 and wild-type or loop-substituted miniGα_s_ (**i**) following PTH1R activation. Luminescence signals were normalized to the response observed with wild-type. **(h**, **j)** Relative expression levels of wild-type and loop-substituted βarr1 (**h**) and miniGα_s_ (**j**), normalized to their respective wild-type controls. **(k)** Structural alignment of the PTH1R ternary complex with the GLP-1R–G_s_ complex (PDB: 7DUQ), superimposed via their Gα_s_ subunits, revealing conserved βarr1-interacting surfaces on Gα_s_. **(l**, **n)** NanoBiT assays of interactions between miniGα_s_ and wild-type or loop-substituted βarr1 (**l**), and between βarr1 and wild-type or loop-substituted miniGα_s_ (**n**) following GLP-1R activation. Luminescence signals were normalized to the response observed with wild-type. **(m, o)** Expression analysis of wild-type and loop-substituted βarr1 (**m**) and miniGα_s_ (**o**) in the GLP-1R system, normalized to their respective wild-type controls. Data are presented as mean ± S.E.M. from at least three independent experiments.

To determine the functional contribution of these three loops, we replaced each with a 5× glycine linker, disrupting specific contacts while preserving the overall protein fold. Disruption of any single node significantly reduced both sustained cAMP accumulation and βarr1**–**Gα_s_ association despite comparable expression (**Fig. 4f–h**, **Extended Data Fig. 1j**, and **Extended Data Table 3**). Reciprocal mutations on the partner Gα_s_ surfaces — Switch I (residues 63**–**66 and 205**–**210), Switch II (residues 232**–**240), and the α3/β5-loop (residues 279**–**285) — similarly impaired the interaction with βarr1 (**Fig. 4i, j**, and **Extended Data Table 4**). Single-point substitutions at individual interfacial sites in miniGα_s_ weakened βarr1 binding to a lesser extent than the corresponding full-loop replacements (**Extended Data Fig. 1k** and **Extended Data Table 5**), underscoring the cooperative, distributed nature of the interface.

These findings establish direct βarr1**–**Gα_s_ coupling as a critical determinant of both complex stability and sustained cAMP signaling. Because the same loop substitutions that break the interface also collapse the internalization-dependent phase of cAMP production, the structural contact and the functional output are linked, and the ternary assembly provides a mechanistic framework for spatiotemporally organized GPCR signaling.

### The βarr1–Gα_s_ interface is conserved across class B GPCRs

We next asked whether this mechanism is a general property of class B GPCRs rather than a feature unique to PTH1R. Dyngo-4a reduced agonist-induced cAMP accumulation across all receptors tested — GLP-1R, GCGR, parathyroid hormone 2 receptor (PTH2R), pituitary adenylate cyclase 1 receptor (PAC1R), and calcitonin receptor (CTR) — without affecting βarr1 recruitment (**Extended Data Fig. 8a, b**). Likewise, overexpression of dominant-negative Dyn2_K44E suppressed cAMP production relative to wild-type Dynamin-2 (**Extended Data Fig. 1c, 8c**). Internalization-dependent Gα_s_ signaling is therefore a conserved, rather than receptor-specific, feature of class B GPCRs.

We then asked whether the structural basis of the βarr1–Gα_s_ interface is similarly conserved. Structural alignment of the PTH1R ternary complex with the GLP-1R–G_s_ complex^49^ showed that the βarr1-binding surface on Gα_s_ is highly conserved (**Fig. 4k**). Consistent with this, NanoBiT assays confirmed that GLP-1R natively forms a ternary complex with βarr1 and miniGα_s_ (**Fig.4l, n**). Introducing the identical βarr1 and miniGα_s_ loop substitutions that disrupted PTH1R assembly produced comparable coupling defects at GLP-1R, at similar expression levels (**Fig. l–o**, and **Extended Data Table 6**). The conserved βarr1–Gα_s_ interface therefore drives complex formation across class B GPCRs, providing a unifying structural basis for βarr-dependent sustained G_s_ signaling.

## Discussion

Endosomal GPCR signaling has long challenged the classical paradigm in which β-arrestin terminates G protein activation by occluding the receptor core. Internalized receptors are known to sustain G_s_/cAMP signaling in a predominantly βarr-dependent manner, but the underlying mechanism has remained unresolved: how can βarr promote rather than terminate G protein activation? Previously described megaplexes established that the two transducers can coexist on a single receptor, yet revealed no contact between them, leaving open whether βarr actively supports G protein signaling or merely accommodates it. We show that βarr1 engages a nucleotide-free state of Gα_s_, through an interface that is required for both complex assembly and the internalization-dependent phase of cAMP production. This interaction is stronger than βarr1–Gβγ coupling and drives cooperative receptor–βarr1–Gα_s_ assemblies that maintain signaling beyond the plasma membrane. Sustained endosomal GPCR signaling therefore rests on direct structural cooperation between β-arrestin and G protein, not merely on their spatial coexistence.

Our cryo-EM structure explains how two canonically opposing transducers can simultaneously occupy one GPCR. Unlike core-engaged complexes that sterically exclude G proteins, βarr1 adopts a tail conformation, anchoring to the receptor C-terminus while leaving the core accessible to Gα_s_. Beyond this spatial accommodation, the defining feature is a direct βarr1**–**Gα_s_ interface where the finger, C-, and back loops of βarr1 engage the Switch I, Switch II, and α3/β5-loop regions of Gα_s_ (**Fig. 4**). This geometry differs from the reported β2V2R and β2AR megaplexes^30,31^, in which βarr1 sits distally with no inter-transducer contact. Structure-guided mutagenesis shows that disrupting these contacts impairs both complex formation and sustained cAMP signaling. Furthermore, this architecture extends beyond PTH1R: blocking internalization suppresses sustained G_s_ activation across multiple class B GPCRs, and the βarr1**–**Gα_s_ interface is functionally conserved at GLP-1R. These contacts therefore define a generalizable mechanism rather than a receptor-specific feature.

Mechanistically, βarr1 associates with the nucleotide-free state of Gα_s_. Specifically, βarr1 associates with two structurally independent nucleotide-free surrogates — the minimized miniGα_s_ and the full-length dominant-negative G112. By holding this intermediate, βarr1 stabilizes Gα_s_ in a signaling-competent state after receptor internalization, thereby prolonging cAMP production. Thus, rather than establishing a direct nucleotide-state preference, our findings identify a structural interaction with a conformation of Gα_s_ that is captured by nucleotide-free surrogate proteins. This interaction could contribute to G protein reactivation after receptor internalization, but the precise role of βarr1 in the nucleotide cycle awaits future direct comparison of different nucleotide-bound states.

However, several features of this work define its current limits. The structure required an engineered assembly — a V2RT chimera, a receptor-βarr1 fusion, scFv30, and miniGα_s_ in place of heterotrimeric G_s_ — thus, the resolved geometry is best regarded as a stabilized snapshot of an inherently transient state. Three observations argue that it nevertheless represents a genuine signaling intermediate rather than an artifact of stabilization. First, the chimeric receptor behaves like the wild-type PTH1R in both NanoBiT and cAMP assays (**Extended Data Fig. 1g**). Second, βarr1**–**Gα_s_ complementation is readily detected in cells using untethered, separately expressed components, and shows the same requirement for a phosphorylated receptor C-tail. Third, mutations designed from the structure produce matched defects in complex formation and in sustained cAMP signaling at two different class B receptors. Importantly, our evidence that this complex operates in the endosomal compartment is functional — it rests on the internalization dependence of the sustained cAMP phase — rather than a direct visualization of the ternary assembly within endosomes; imaging the complex in that compartment remains an important next step.

Integrating these results with a previously proposed PTH1R**–**βarr**–**Gβγ complex^29^, we propose a model of dynamic G protein cycling on the internalized receptor (**Fig. 5**). In that earlier model, βarr promotes G_s_ activation by stabilizing a PTH1R**–**Gβγ complex; our present structure supplies a candidate mechanistic step for Gα_s_. Following GTP hydrolysis, GDP-bound Gα_s_ reassociates with the receptor**–**βarr1 scaffold, and βarr1 stabilizes the nucleotide-free intermediate, potentially facilitating a new cycle of nucleotide exchange. The βarr1**–**Gβγ and βarr1**–**Gα_s_ assemblies would then represent sequential states of a reactivation pathway where βarr1 serves as the organizational scaffold. Despite extensive cryo-EM effort, we did not capture a stable PTH1R**–**βarr1**–** Gα_s_βγ or PTH1R**–**βarr1**–**Gβγ complex, consistent with the spatial separation or high conformational heterogeneity of the G protein subunits. Defining how Gβγ is coordinated within this framework therefore remains a critical next step.

**Fig. 5.**
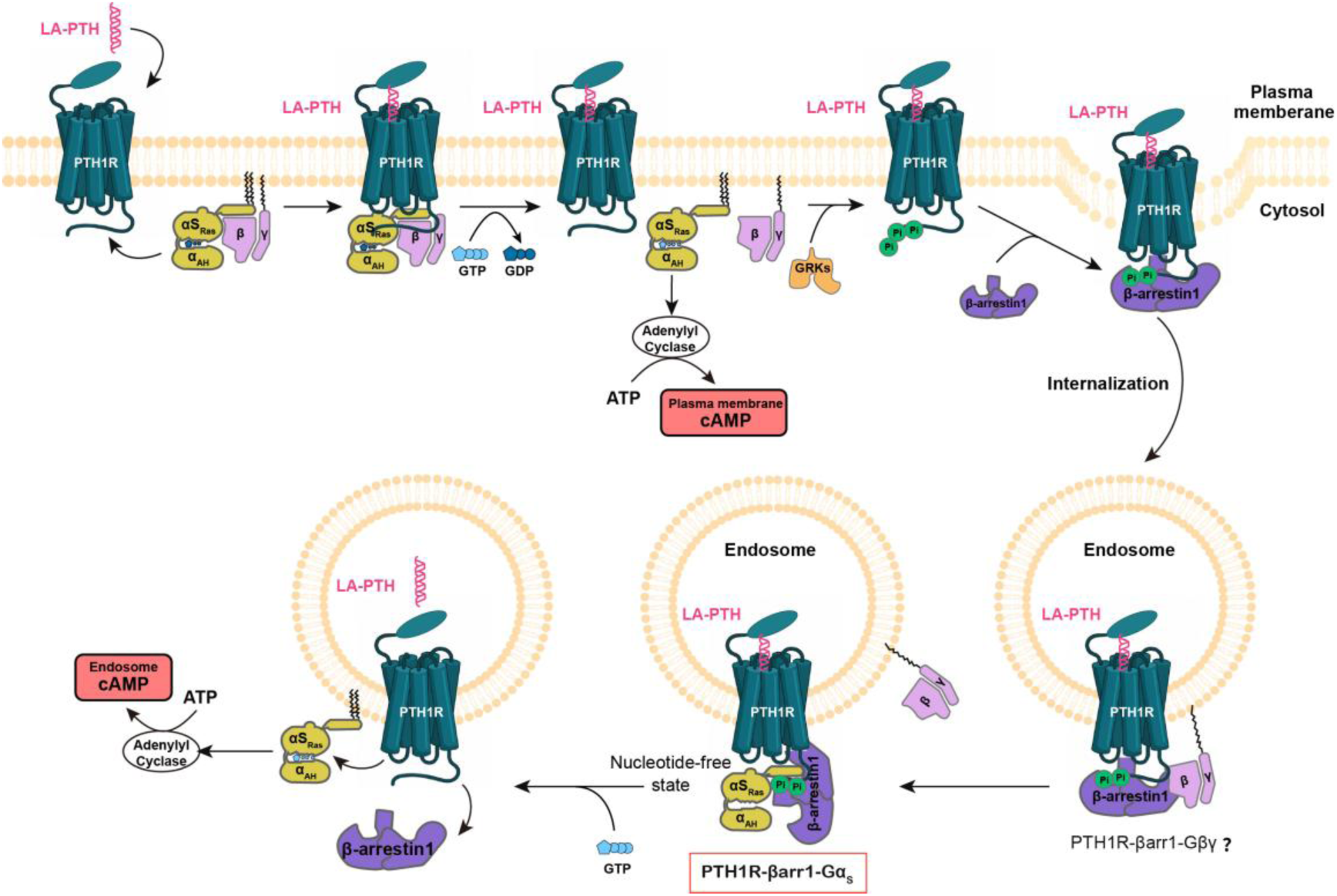
Proposed pathway of PTH1R-βarr1-Gα_s_ complex assembly and sustained cAMP signaling. Production of cAMP (1^st^ pool) by the PTH-bound receptor at the plasma membrane is short-lived due to receptor desensitization by GPCR kinases (GRKs) and β-arrestin (βarr) binding to the fully phosphorylated (P) receptor. The PTH-bound PTH1R in complex with βarr internalizes and redistributes to endosomes. We propose that βarr stabilizes active Gα_s_ in endosomes by assembling the complex, supporting sustained cAMP production (2^nd^ pool) for PTH signaling.

Ultimately, these findings revise the classical binary model of GPCR regulation. Rather than simply handing off from G protein activation to arrestin-mediated desensitization, internalized receptors assemble cooperative complexes in which βarr1 and Gα_s_ act together to sustain cAMP signaling, with the direct βarr1**–**Gα_s_ interface providing the structural basis for that cooperation. Beyond establishing a framework for spatially organized GPCR signaling, these findings provide a molecular rationale for the prolonged pharmacological actions of PTH1R-targeted therapies; the investigational agonist eneboparatide, for example, stabilizes a prolonged-signaling receptor conformation to sustain calcemic responses. By defining the interface that generates this sustained signal, our findings offer a mechanistic foundation for designing agonists with tailored signaling duration and compartmental selectivity.

## Materials and methods

### Plasmid construction

The human PTH1R (residues 27–502) fused with the V2R C-terminal tail (V2RT, residues 343–371^9,43^) was cloned into the pFastBac vector (Invitrogen), preceded by an N-terminal hemagglutinin (HA) signal peptide and thermostabilized BRIL. A modified β-arrestin1 (residues 3–393, containing 3A mutations: I386A, V387A, F388A^50^) was fused to the C-terminus of PTH1R via a GSA linker, followed by single-chain Fab30 (scFv30) for complex stabilization^4,44,51^, and a C-terminal 8× His-tag. Wild-type bovine GRK2 and engineered miniGα_s_ were also cloned into pFastBac. The miniGα_s_ construct was truncated to remove the α-helical domain (AHD; G67–T204) and residues N254– T263, and included stabilizing mutations (G49D, E50N, L63Y, A249D, S252D, L272D, A366S, I372A and V375I).

For functional assays, wild-type and mutant PTH1R and other GPCRs were cloned into pcDNA6.0 (Promega). Constructs of miniGα_s_, G112, Gβ1, and Gγ2 were cloned into pBiT2.1 (Promega) with the LgBiT subunit at the N-/C-terminus for NanoBiT complementation assays. For G112 — a modified bovine Gα_s_ (mDNGα_s_) as previously reported^42^ — the N terminus (M1–K25) and α-helical domain (AHD F68–L203) of Gα_s_ were replaced with the N terminus (M1-M18) and AHD (Y61–K180) of human Gα_i_, residues N254–T263 of Gα_s_ were deleted, and eight dominant-negative mutations (G49D, E50N, L63Y, A249D, S252D, L272D, I372A and V375I) were incorporated. The same βarr1 construct was cloned into pBiT2.1 with SmBiT at the N-terminus, enabling luminescent signal generation upon interaction with LgBiT-tagged partners.

### Expression of PTH1R–βarr1–miniGα_s_ ternary complex

To promote formation of the ternary complex, PTH1R_V2RT–βarr1–scFv30, miniGα_s_, and GRK2 were co-expressed in Sf9 insect cells (Invitrogen) at a density of 4.0×10^6^ cells/mL in serum-free SIM SF Expression Medium (SinoBiological). Sf9 cells were infected with recombinant baculoviruses encoding the three components at the ratio of 2:2:1 and cultured at 27 ℃ for 48 hours. Cells were harvested by centrifugation, resuspended in PBS, and centrifuged again. The resulting cell pellets were flash-frozen and stored at −80 ℃ until further use.

### Purification of PTH1R‒βarr1‒miniGα_s_ ternary complex

Frozen cell pellets were thawed at room temperature and resuspended in lysis buffer containing 20 mM HEPES (pH 7.40), 100 mM NaCl, 10 % glycerol, 5 mM MgCl_2_, 1 mM ATP, 2 mM Na_3_VO_4_, 100 μM TCEP, and 1× Protease Inhibitor Cocktail (TargetMol). To promote receptor phosphorylation and complex formation, 10 μM LA-PTH was added, and the suspension was incubated at room temperature for 1 hour. The lysate was then clarified by centrifugation at 30,000 rpm for 30 minutes. The resulting pellet was resuspended in solubilization buffer consisting of 20 mM HEPES (pH 7.40), 100 mM NaCl, 10 % glycerol, 5 mM MgCl_2_, 100 μM TCEP, 1× Protease Inhibitor Cocktail, and 10 μM LA-PTH, and solubilized with 0.5 % (w/v) LMNG and 0.1 % (w/v) CHS at 4 ℃ for 3 hours.

Following solubilization, the supernatant was collected by centrifugation at 30,000 rpm for 45 minutes and incubated with pre-equilibrated Ni Focurose FF (IDA) resin in the presence of 20 mM imidazole at 4 ℃ for 1.5 hours. The resin was then loaded into a gravity-flow column and washed sequentially with 15 column volumes of two buffers: first with 20 mM HEPES (pH 7.40), 100 mM NaCl, 10 % glycerol, 2 mM MgCl_2_, 25 μM TCEP, 0.01 % LMNG, 0.01 % GDN, 0.004 % CHS, 2 μM LA-PTH, and 30 mM imidazole, followed by the same buffer containing 45 mM imidazole to remove nonspecific binders. Elution was performed using 10 column volumes of elution buffer containing 300 mM imidazole. The eluate was concentrated using a 100-kDa Amicon Ultra Centrifugal Filter (Millipore).

For complex stabilization, the sample was incubated with Nb32 at a 1:10 molar ratio for 2 hours on ice. The complex was further purified by size-exclusion chromatography using a Superose 6 Increase 10/300 GL column (GE Healthcare) equilibrated with buffer containing 20 mM HEPES (pH 7.40), 100 mM NaCl, 2 mM MgCl_2_, 25 μM TCEP, 0.00075 % LMNG, 0.00025 % GDN, 0.0001 % digitonin, 0.0002 % CHS, and 2 μM LA-PTH. Fractions containing the target complex were collected and concentrated for cryo-EM grid preparation.

### Expression and purification of Nb32

Nb32 was cloned into the pMESy4 vector with a C-terminal 6×His-tag and expressed in E. coli BL21 cells. Cells were grown in TB medium supplemented with 100 mg/mL ampicillin, 2 mM MgCl_2_, and 0.1 % glucose at 37 ℃, 190 rpm for approximately 5 hours. Upon reaching an OD_600_ of ∼1.0, protein expression was induced with 0.5 mM IPTG and continued overnight at 20 ℃, 190 rpm.

Cells were harvested and lysed using a high-pressure homogenizer in lysis buffer containing 20 mM HEPES (pH 7.40), 100 mM NaCl, and 1 mM PMSF at 4 ℃. The lysate was clarified by centrifugation at 30,000 rpm for 30 minutes, and the supernatant was incubated with pre-equilibrated Ni Focurose FF (IDA) resin in the presence of 10 mM imidazole at 4 ℃ for 1 hour. The resin was loaded into a gravity-flow column and washed with 30 column volumes of wash buffer containing 20 mM HEPES (pH 7.40), 100 mM NaCl, and 25 mM imidazole. Protein was eluted with 10 column volumes of elution buffer (20 mM HEPES, pH 7.40; 100 mM NaCl; 300 mM imidazole) and concentrated using a 10-kDa Amicon Ultra centrifugal filter (Millipore). The sample was further purified on a HiLoad 16/600 Superdex 75 column equilibrated with 20 mM HEPES (pH 7.40) and 100 mM NaCl. The final purified Nb32 was concentrated and stored at −80 ℃.

### NanoBiT complementation assay

The interaction between miniGα_s_/G112/Gβ1/Gγ2 and βarr1 upon receptors activation was measured using NanoLuc^®^ Binary Technology (NanoBiT^®^, Promega) in HEK293 cells. Cells were seeded into 12-well or 6-well plates and cultured in DMEM (high glucose, Cytiva) supplemented with 10 % (v/v) fetal bovine serum at 37 ℃ in 5 % CO_2_. After approximately 20 hours, cells were transfected using PEI MAX 40K with plasmids encoding pcDNA-PTH1R/GLP-1R, pBiT-LgBiT–Gα_s_/Gβ1/Gγ2 (alternatively, pBiT-miniGα_s_/Gβ1/Gγ2–LgBiT), and pBiT-SmBiT–βarr1, at a 1:1:1 ratio.

Following a 24-hour transfection, cells were harvested by centrifugation at 100 × g for 10 minutes, resuspended in PBS, and dispensed into 384-well plates at 10 μL per well, with a final cell density of 5.0 × 10^5^ cells/mL. Coelenterazine 400a (diluted in PBS) was added to each well at 5 μL per well to a final concentration of 10 μM in the dark. Subsequently, LA-PTH at varying concentrations was added (5 μL per well), and luminescence signals were recorded using an EnVision plate reader (PerkinElmer) at a speed of 1 s per well.

For mutant constructs, miniGα_s_ or βarr1 variants were cloned into the pBiT2.1 vector and transfected using the same procedure. To assess the influence of Gβγ or miniGα_s_, the corresponding genes were cloned into pcDNA3.1 vector and co-transfected with the constructs mentioned above at equal ratios. The empty pcDNA3.1-Vector was used as a negative control.

The interaction between receptors and βarr1 upon receptor activation was also measured using NanoLuc^®^ Binary Technology (NanoBiT^®^, Promega). The LgBiT subunit was fused to the C-terminus of receptors and the SmBiT subunit was fused to the N-terminus of βarr1. Both constructs were cloned to the pBiT2.1 vector. To assay the effect of dynamin inhibitor Dyngo-4a in the βarr1 recruitment, Dyngo-4a was co-applied with agonists and added to 384-well plates at the final concentrations of 20 μM. DMSO, used as the solvent for Dyngo-4a, was added with agonists as the vehicle control. Data were analyzed using GraphPad Prism 9.5 and are presented as mean ± S.E.M. from at least three independent experiments.

### GloSensor cAMP accumulation assay

Intracellular cAMP production was assessed using the GloSensor^TM^ cAMP Assay (Promega) in HEK293 cells. Cells were seeded in 6-well plates and cultured overnight in DMEM (high glucose, Cytiva) supplemented with 10 % (v/v) fetal bovine serum at 37 ℃, 5 % CO_2_. The next day, pcDNA-receptors and GloSensor-22F plasmids were co-transfected at a 1:1 ratio using PEI MAX 40K. After ∼20 hours, cells were collected and seeded into 96-well plates at 100 μL/well at a density of 3.0×10^5^ cells/mL.

After 18 hours of growth in medium, the DMEM was aspirated and replaced with CO_2_-independent medium containing 2 % GloSensor cAMP Reagent at 50 μL/well, followed by a 1-hour incubation at 37 ℃. Ligands were prepared in CO_2_-independent medium and added directly to each well. Luminescence signals were recorded immediately using a Synergy H1 microplate reader (BioTek) at 30 seconds per plate.

To assess the effect of βarr1, HEK293 cells were transfected one day prior to plasmid transfection with either a negative control siRNA or an siRNA specifically designed to target the βarr1 sequence in the exogenous βarr1 construct used in this study. On the following day, cells were co-transfected with pcDNA-PTH1R, GloSensor-22F, and either pcDNA3.1-βarr1 or pcDNA3.1-Vector at a 1:1:1 molar ratio. pcDNA3.1-βarr1 was transfected into both βarr1-specific siRNA- and control siRNA-treated cells, while pcDNA3.1-Vector was introduced only into the control cells.

To evaluate endosomal cAMP signaling, the dynamin inhibitor Dyngo-4a was mixed with agonists and added to 96-well plates at the final concentrations of 20 μM or 50 μM. In parallel, cells were co-transfected with pcDNA-PTH1R and GloSensor-22F along with either wild-type Dynamin2 or dominant-negative Dynamin2 mutant K44E to evaluate the effect of impaired dynamin function.

In a separate approach, cells were first stimulated with LA-PTH for 15 minutes, followed by washing twice with 100 μL CO_2_-independent medium. Subsequently, 50 μL/well of CO_2_-independent medium containing 2 % GloSensor cAMP Reagent was added, and cells were incubated at 37 ℃ for 1 hour prior to luminescence detection. Data were analyzed using GraphPad Prism 9.5 and are presented as mean ± S.E.M. from at least three independent experiments.

### Detection of expression levels

Expression levels of miniGα_s_ or β-arrestin1 were assessed by flow cytometry using an N-terminal 3× FLAG (DDDDK) epitope tag. HEK293 cells were harvested 24 h post-transfection and washed once with 1 % BSA (w/v, in 1× PBS). To allow antibody access to intracellular epitopes while preserving overall cell morphology, cells were gently permeabilized with 0.01 % digitonin (prepared in 1× PBS) for 3 min at room temperature (RT), followed by two washes with 1× PBS. Cells were then blocked with 5 % BSA (w/v, in 1× PBS) for 30 min at RT and incubated with a mouse anti-DDDDK monoclonal antibody (ABclonal; 1:100 dilution in 5 % BSA) for 30 min at RT. After three washes with 1 % BSA, cells were incubated on ice and protected from light with an ABflo^®^ 488-conjugated goat anti-mouse IgG (H+L) secondary antibody (ABclonal; 1:1000 dilution in 5 % BSA) for 1 h. Cells were subsequently washed three times with 1 % BSA, resuspended in 1 % BSA, and analyzed on Guava EasyCyt^TM^ flow cytometer (excitation: 488 nm; emission: 525 nm), acquiring approximately 5,000 events per sample. Data were analyzed using GuavaSoft^TM^ software (version 4.5), normalized to wild-type controls, and presented as mean ± S.E.M. from at least three independent experiments.

### Cryo-EM grid preparation and data collection

A volume of 3 μL purified LA-PTH–PTH1R–βarr1–miniGα_s_ complex at a concentration of 11.65 mg/mL was applied to glow-discharged UltrAuFoil 300 mesh grids (Quantifoil R1.2/1.3, Au/Au or Au/C). Grids were plunge-frozen in liquid ethane using a Vitrobot Mark VI (Thermo Fisher Scientific) with a 5-second wait time, blot force of 2, and 4-second blot time, under 100 % humidity at 4 ℃. Cryo-EM data were collected on a Titan Krios G4 electron microscope (Thermo Fisher Scientific) operated at 300 kV, equipped with a Falcon4i direct electron detector and Selectris X energy filter. Images were recorded using EPU software (Thermo Fisher Scientific) in electron event registration (EER) mode at a physical pixel size of 0.73 Å. A total electron dose of 50 e^−^/Å^2^ was used at a dose rate of 10 e^−^ pixel^−1^ s^−1^, and data were collected with a defocus range of −0.8 to −1.8 μm.

### Cryo-EM images processing

A total of 111,056 cryo-EM movies of the LA-PTH‒PTH1R‒βarr1‒miniGα_s_ complex were motion-corrected using RELION^52^, and the contrast transfer functions (CTFs) were estimated with the Patch CTF estimation module in CryoSPARC^53^. Micrographs were curated based on CTF fit resolution and astigmatism. Particle picking was performed using Blob Picker, Template Picker, and Topaz (models from a partial dataset). The picked particles were extracted with a box size of 360 pixels (binned by 4) and classified in multiple rounds of 2D classification and heterogeneous refinement using initial models generated by ab-initio reconstruction. The best-performing particles were re-extracted with a 440-pixel box size and subjected to heterogeneous refinement using initial models generated by ab-initio reconstruction, yielding six classes. Final particle stacks of 194,863 particles were processed via non-uniform refinement.

To improve local resolution, various subregions of the complex — including PTH1R, miniGα_s_, βarr1, scFv30, and critical interfaces — were individually refined using mask-based local refinement. The composite map was assembled by aligning the locally refined maps to the consensus map and combining selected maps using the maximum value of each component (vop maximum) in UCSF ChimeraX^54,55^. In addition, DeepEMhancer^56^ was applied to enhance map features from an unsharpened composite map.

### Model building and refinement

Locally refined and composite cryo-EM maps were used for model building unless otherwise specified. All maps were sharpened using the B-factor automatically calculated in CryoSPARC and DeepEMhancer. AlphaFold2^57^ predicted models of the protein complex (PTH1R, miniGα_s_, β-arrestin1, and scFv30) were used as templates and docked into the cryo-EM maps from DeepEMhancer. The models were iteratively refined using Coot^58^ and Phenix^59^ Real Space Refinement using B-factor cryo-EM maps. The final refinement was validated using Phenix.

### Quantification and statistical analysis

All cell-based assays were performed in at least three independent experiments, each in triplicate, and are presented as mean ± S.E.M. unless stated otherwise. Concentration-response data from NanoBiT complementation and GloSensor cAMP assays were fitted with a three-parameter logistic model in GraphPad Prism 9.5 to obtain pEC_50_ and span values. Comparisons among wild-type and mutant constructs were made by two-sided one-way ANOVA with Tukey’s multiple comparisons test; *P < 0.05, **P < 0.01, ***P < 0.001, ****P < 0.0001. Expression levels of all mutants were measured in parallel by flow cytometry and used to confirm that changes in signal were not attributable to differences in protein abundance. No data were excluded from the analyses, and no statistical method was used to predetermine sample size.

## Data availability

The cryo-EM density map and atomic coordinates of the LA-PTH-bound PTH1R–β-arrestin 1–Gα_s_ ternary complex have been deposited in the Electron Microscopy Data Bank (EMDB) and Protein Data Bank (PDB) under accession codes EMD-69688 and 24NJ, respectively. The corresponding split and masked cryo-EM maps have been deposited under EMDB accession codes EMD-69689 (mask on Gα_s_), EMD-69690 (mask on β-arrestin 1 and scFv30), EMD-69691 (mask on β-arrestin 1 and scFv30), EMD-69692 (mask on Gα_s_ and β-arrestin 1), EMD-69693 (mask on PTH1R–Gα_s_), and EMD-69694 (LA-PTH-bound PTH1R–β-arrestin 1–Gα_s_ ternary complex). Previously published structures used for comparison are available under PDB accession codes 6NBH, 8JRV, 6U1N, 8WU1, 7R0C, 6NI2, 6NI3, 9L8L, 7DUQ and 1AZT. All other data supporting the findings of this study are available within the article and its Supplementary Information, or from the corresponding authors upon reasonable request.

## Acknowledgements

We thank the Advanced Center for Electron Microscopy at Shanghai Institute of Materia Medica, Chinese Academy of Sciences, where cryo-EM data were collected. This work was supported by the National Natural Science Foundation of China (32371255 and 32071203 to L.H.Z., 32130022 and 82121005 to H.E.X., 82404881 to Q.N.Y.); the Natural Science Foundation of Shanghai (23ZR1475200 to L.H.Z.); the National Key R&D Program of China (2022YFC2703105 to H.E.X., 2019YFA0904200); the CAS Strategic Priority Research Program (XDB37030103 to H.E.X.); the Shanghai Municipal Science and Technology Major Project (2019SHZDZX02 to H.E.X.); the Young Innovator Association of CAS (Y2022078 to L.H.Z.); the Lingang Laboratory (LG-GG-202204-01 to H.E.X.); and the State Key Laboratory of Drug Research (SKLDR-2023-TT-04 to H.E.X.).

## Declaration of generative AI and AI-assisted technologies in the writing process

During the preparation of this work, we used AI-assisted tools, including Claude.AI, Super Grok, and ChatGPT, to improve language refinement and writing clarity. All content generated with these tools was carefully reviewed and edited by the authors, who take full responsibility for the final version of the manuscript.

## Author contributions

L.H.Z. and H.E.X. initiated the investigation into the function and structure of the parathyroid hormone 1 receptor (PTH1R) ternary complex, organized the entire project, and supervised the overall experimental design and execution. L.H.Z. and H.E.X. participated in data analysis and interpretation. Q.H. designed the expression and functional constructs, purified the protein complexes, prepared samples for cryo-EM data collection, conducted the NanoBiT, cAMP and flow cytometry assays, analyzed structure and function data, prepared figures, and drafted the manuscript. Q.N.Y. performed cryo-EM map calculations, and built and refined the structural models. X.H. performed structural analysis and comparison, and contributed to figure preparation. W.H. collected the cryo-EM data. L.H.Z. provided the original plasmid for protein purification. L.H.Z. and H.E.X. wrote the manuscript with input from all the authors.

## Competing interests

All authors declare that they have no competing interests.

## Notes

### Competing Interest Statement

The authors have declared no competing interest.

