## Extended Data Fig. 1 to 8 and Extended Data Table 1 to 6 for "β-arrestin1 directly engages Gαs to sustain endosomal GPCR signaling"

### Supplementary Information

**Extended Data Fig. 1 to 8**

**Extended Data Table 1 to 6**

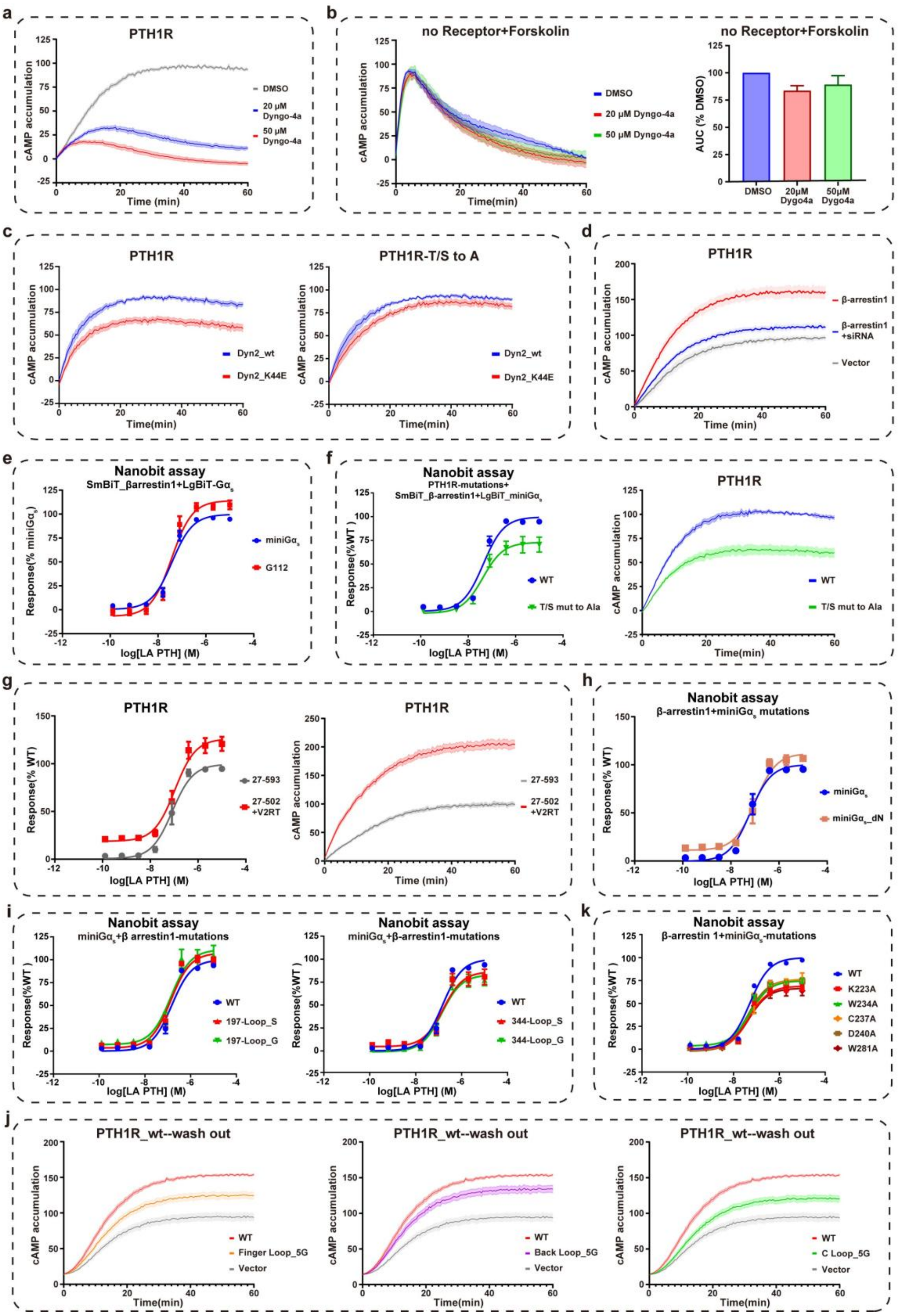


**Extended Data Fig. 1 GloSensor cAMP accumulation and NanoBiT complementation assays of PTH1R.**

**(a)** Dyngo-4a inhibition of LA-PTH-stimulated cAMP accumulation in HEK293 cells expressing PTH1R_WT. Luminescence signals were normalized to vehicle-treated control (DMSO).

**(b)** Effect of Dyngo-4a on forskolin-stimulated cAMP in cells expressing the GloSensor biosensor alone. Luminescence was recorded over 60 min and normalized to DMSO-treated controls (**left**). Area under the curve (AUC) values were calculated and normalized accordingly (**right**).

**(c)** Effects of wild-type or dominant-negative dynamin-2 (Dyn2_K44E) on cAMP accumulation mediated by PTH1R_WT (**left**) or by the mutant with all Ser/Thr residues in the C-tail substituted with Ala (T/S mut to A) (**right**). Luminescence signals were normalized, within each receptor background, to the response observed with Dyn2_WT.

**(d)** Effects of βarr1 overexpression, alone or combined with an siRNA targeting the βarr1 sequence used in this study, on LA-PTH-induced cAMP accumulation. Luminescence was recorded over a 60-minute period and normalized to vector-transfected control.

**(e)** NanoBiT complementation assays measuring interactions between βarr1 and miniG$\alpha_s$ or G112. Luminescence signals were normalized to the response obtained with LgBiT–miniG$\alpha_s$ and SmBiT–βarr1.

**(f)** Comparison of βarr1–miniG$\alpha_s$ interaction (**left**) and cAMP signaling (**right**) between PTH1R_WT and the mutant with all Ser/Thr residues in the C-tail substituted with Ala (T/S mut to A). The signals were normalized to those obtained with PTH1R_WT.

**(g)** Comparison of βarr1–miniG$\alpha_s$ interaction (**left**) and cAMP signaling (**right**) between full-length PTH1R(27–593) and the truncated PTH1R(27–502)_V2RT chimera. The signals were normalized to those observed with full-length PTH1R(27–593).

**(h)** NanoBiT assay of βarr1 interaction with wild-type or N-terminally truncated miniG$\alpha_s$ (miniG$\alpha_s$_dN, lacking the first 37 amino acids). Luminescence signals were normalized to those observed with miniG$\alpha_s$_WT.

**(i)** NanoBiT analysis of the interaction between miniG$\alpha_s$ and βarr1 mutants in which all residues of the 197- (**left**)or 344-loops (**right**) were substituted with Gly or Ser. Luminescence signals were normalized to the response observed with βarr1_WT.

**(j)** cAMP accumulation following ligand washout in cells expressing wild-type βarr1 or loop-substitution mutants. Related to **Fig. 4f**.

**(k)** NanoBiT dose-response analysis of βarr1 interaction with wild-type or single-point mutants of miniG$\alpha_s$ at the βarr1–miniG$\alpha_s$ interface. Luminescence signals were normalized to those observed with miniG$\alpha_s$_WT. Related to **Fig. 4i**.

Data are presented as mean ± S.E.M. from at least three independent experiments, each performed in triplicate.

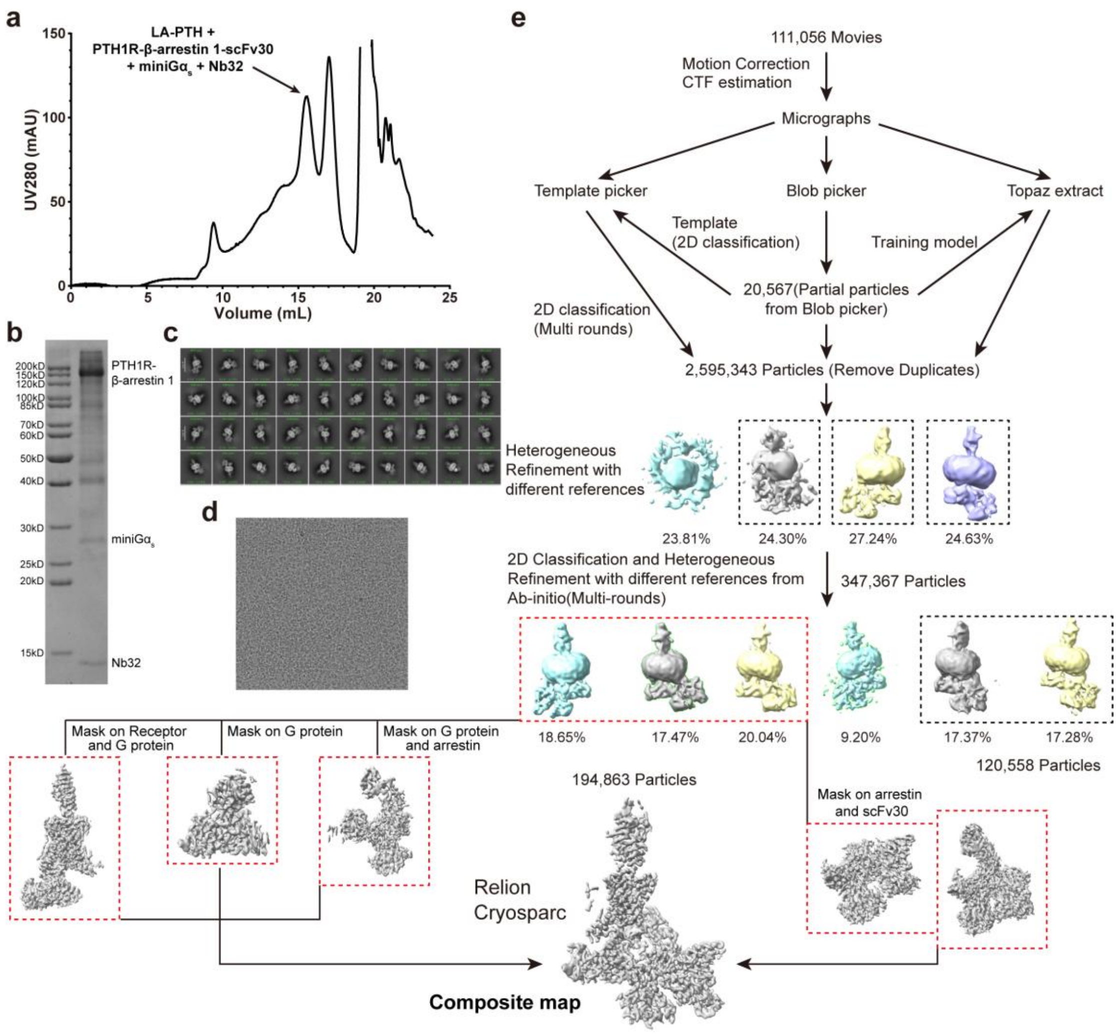


**Extended Data Fig. 2 LA-PTH–PTH1R–βarr1–G$\alpha_s$ complex purification and cryo-EM data processing (related to Fig. 2).**

**(a, b)** Size-exclusion chromatography profile (**a**) and SDS-PAGE analysis (**b**) of the purified PTH1R–βarr1–miniG$\alpha_s$ complex. The black arrow indicates the peak corresponding to the complex monomer.

**(c, d)** Representative 2D class averages (**c**) and cryo-EM micrograph (**d**) of the complex particles.

**(e)** Workflow of cryo-EM single-particle analysis for the PTH1R–βarr1–miniG$\alpha_s$ complex. The final composite map was generated by merging locally refined sub-maps masked on PTH1R–miniG$\alpha_s$, miniG$\alpha_s$ alone, miniG$\alpha_s$–βarr1, and βarr1–scFv30 regions.

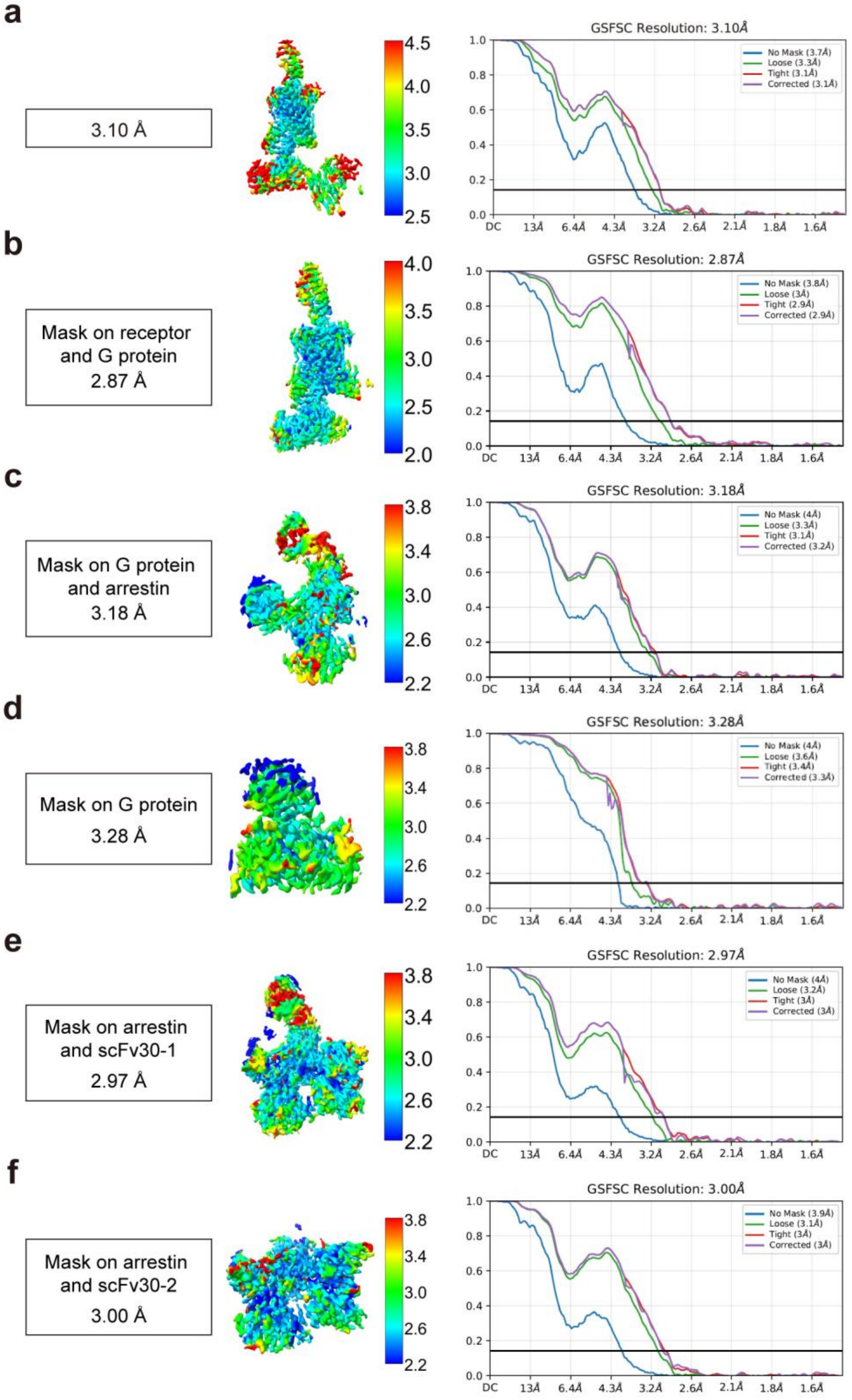


**Extended Data Fig. 3 Color Cryo-EM densities and Fourier Shell Correlation (FSC) curves of the LA-PTH–PTH1R–βarr1–Gα$_s$ complex.**

**(a)** Overall color-coded cryo-EM density map (left) and FSC curves (right) for the calculated map of the PTH1R–βarr1–miniGα$_s$ complex.

**(b)** Locally refined density map (left) and FSC curves (right) for the region masked on the PTH1R–miniG$\alpha_s$ subcomplex.

**(c)** Locally refined density map (left) and FSC curves (right) for the miniG$\alpha_s$–βarr1 interface.

**(d)** Locally refined density map (left) and FSC curves (right) for the miniG$\alpha_s$ region alone.

**(e**, **f)** Locally refined density maps (left) and FSC curves (right) for the βarr1–scFv30 subcomplexes.

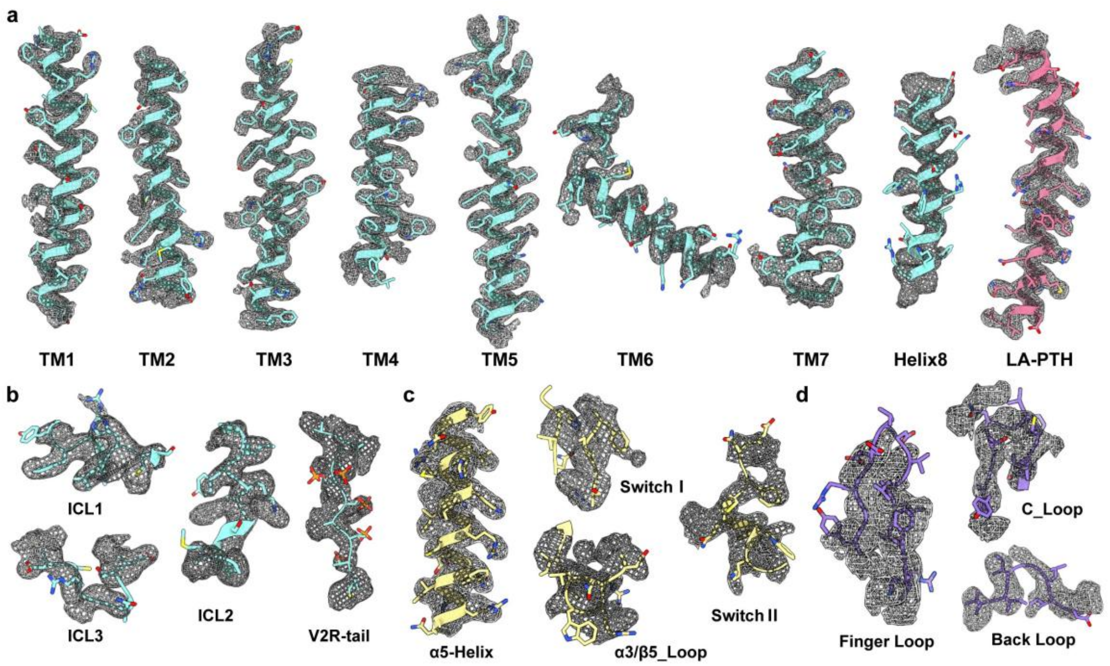


**Extended Data Fig. 4 Density maps of the LA-PTH–PTH1R–βarr1–G$\alpha_s$ complex. Related to Fig. 2.**

**(a, b)** Representative cryo-EM density maps highlighting key regions of PTH1R in the complex: TM1-TM7, helix 8, and bound LA-PTH peptide (**a**); ICL1-ICL3 and the engineered V2R C-terminal tail (V2R-tail) (**b**).

**(c)** Representative cryo-EM densities of miniG$\alpha_s$ showing the main features for the α5-helix, Switch I, Switch II, and the α3/β5 loops.

**(d)** Cryo-EM density map of βarr1 displaying significant structural elements including the finger loop, C-loop, and back loop.

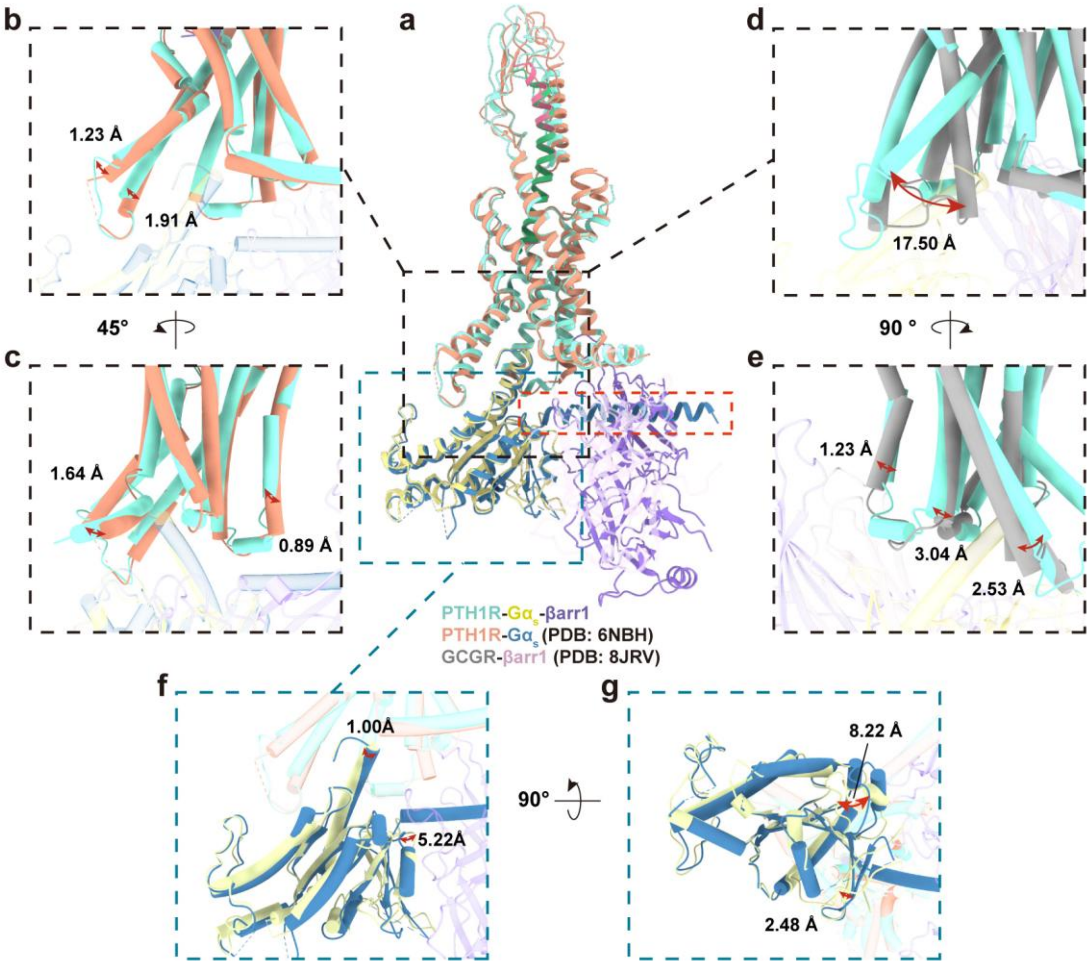


**Extended Data Fig. 5 Distinct conformational features of $G\alpha_s$ and PTH1R.**

**(a)** Structural alignment of the PTH1R–βarr1–miniG$\alpha_s$ complex with the PTH1R–G$\alpha_s$ complex (PDB: 6NBH) and the GCGR–βarr1 complex (PDB: 8JRV), using receptors as the common reference framework.

**(b, c)** Comparison of intracellular conformational changes in PTH1R between the PTH1R–βarr1–miniG$\alpha_s$ ternary assembly and the PTH1R–G$\alpha_s$ complex (PDB: 6NBH), highlighting the displacements of TM4-TM6 and helix 8. Two side views are shown, rotated by 45° along the y-axis.

**(d, e)** Structural overlay of PTH1R from the ternary assembly and GCGR from the GCGR–βarr1(PDB: 8JRV), illustrating rearrangements in intracellular transmembrane helices to enable dual transducers (G protein and βarr1) engagement. Two side views are shown, rotated by 90° along the y-axis.

**(f, g)** Comparison of $G\alpha_s$ conformations between the ternary assembly and the PTH1R–$G\alpha_s$ (PDB: 6NBH), displayed from the side view and the cytoplasmic side, rotated by 90° along the x-axis.

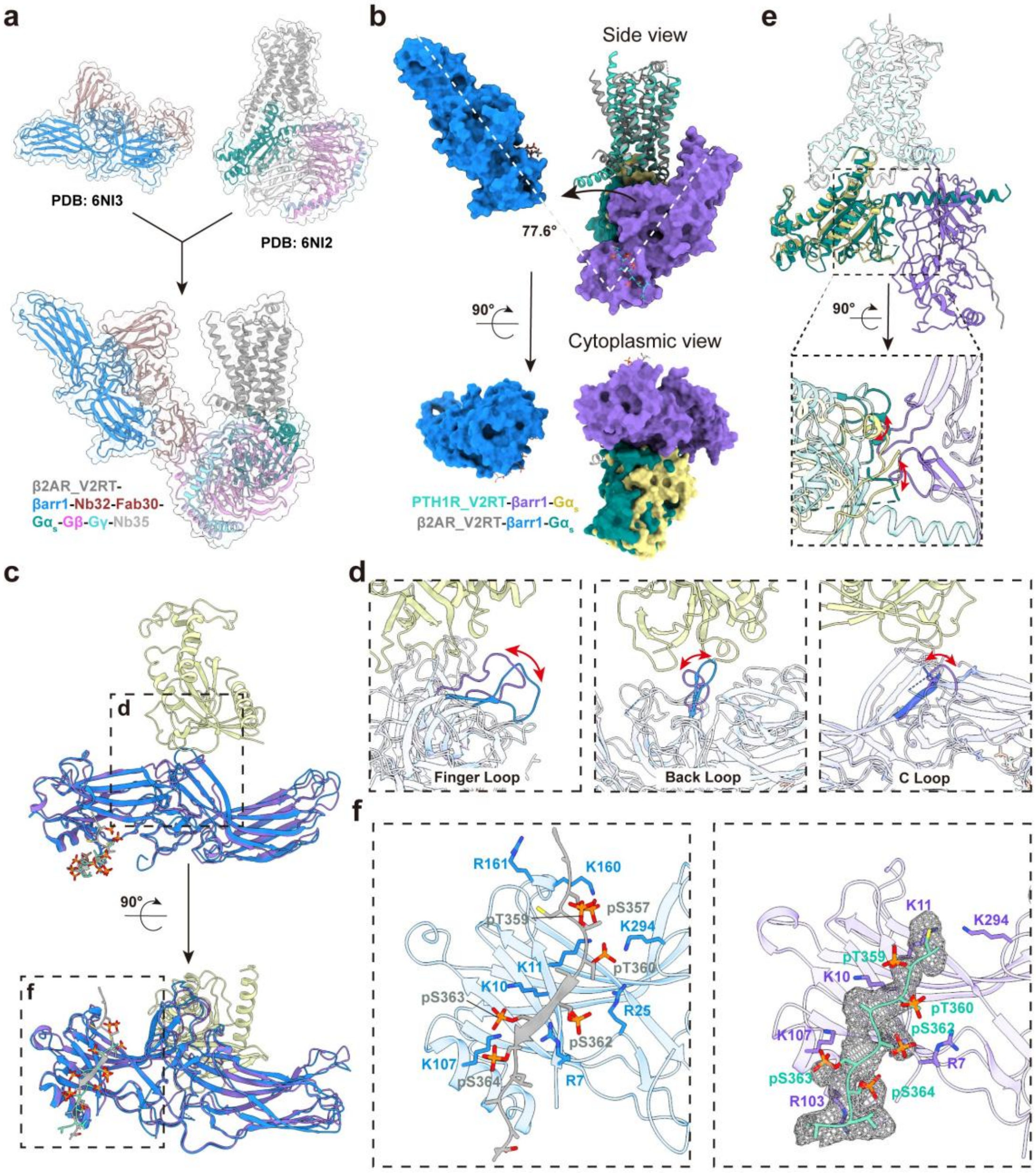


**Extended Data Fig. 6 Structural comparison of the PTH1R–βarr1–miniGα$_s$ with the β2V2R–βarr1–G$_s$ complexes.**

**(a)** Cryo-EM maps of the β2V2R–βarr1–G$_s$ megaplex showing the split maps (V2RT–βarr1–Nb32–Fab30, PDB: 6NI3; β2V2R–G$_s$–Nb35, PDB: 6NI2) and merged map for reference.

**(b)** Structural comparison of βarr1 in the PTH1R–βarr1–miniGα$_s$ and β2V2R–βarr1–G$_s$ complexes, aligned by the receptor transmembrane domains. Upper panels show side views; lower panels show cytoplasmic views rotated 90° along the x-axis.

**(c)** Comparison of the βarr1–Gα$_s$ interface between the two complexes, aligned by βarr1.

The upper panels show side view, and the lower panels show view rotated by 90° along the x-axis.

**(d)** Detailed comparison of the finger loop, back loop, and C-loop regions of βarr1 between the two complexes, aligned by βarr1.

**(e)** Structural comparison of $G\alpha_s$ between the PTH1R–βarr1–mini$G\alpha_s$ and β2V2R–βarr1–$G_s$ complexes, aligned by the receptor transmembrane domains. Upper panels show side views; lower panels show cytoplasmic views rotated 90° along the x-axis.

**(f)** Structural comparison of the receptor C-terminal tails: the V2R tail in β2V2R–βarr1–$G_s$ megaplex (left) and PTH1R–βarr1–mini$G\alpha_s$ complex (right), aligned by βarr1.

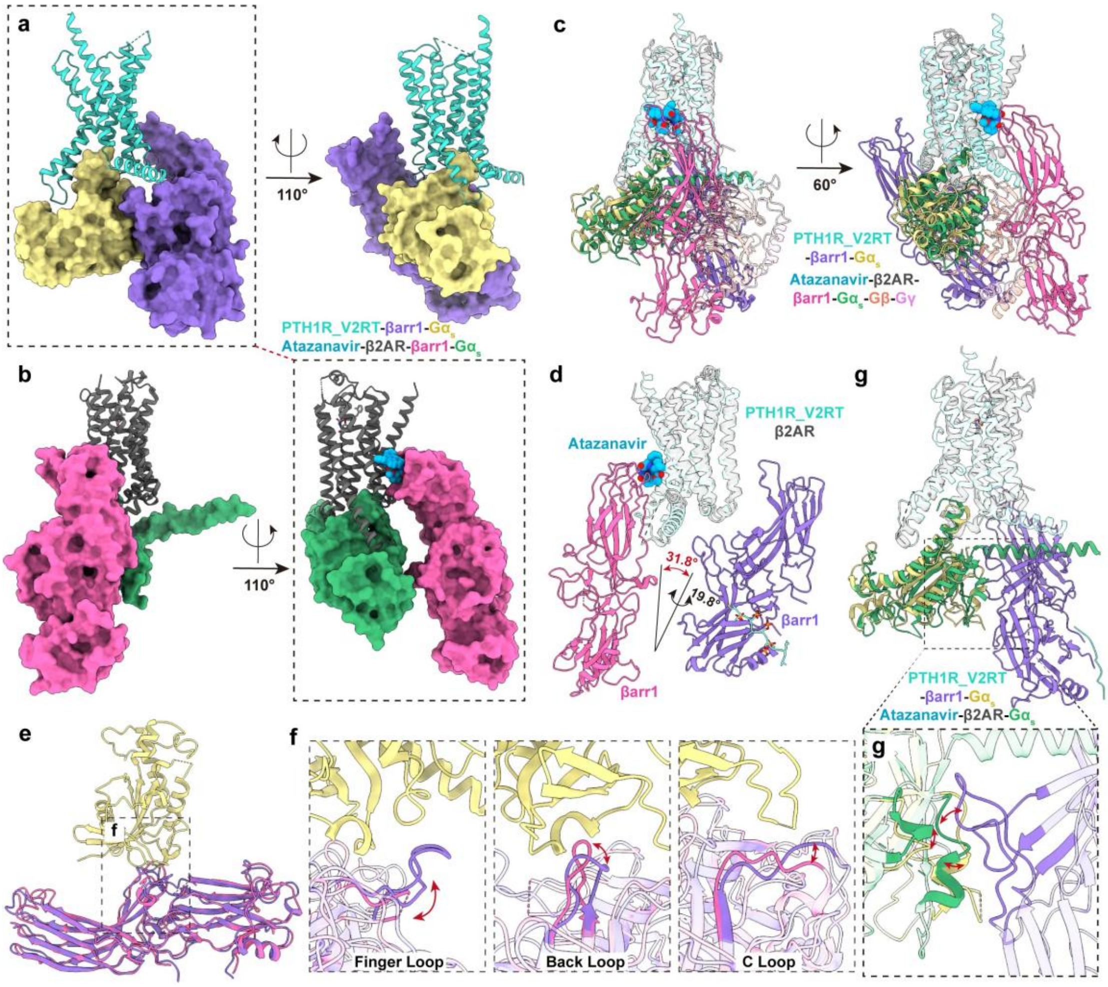


**Extended Data Fig. 7 Structural comparison of the PTH1R–βarr1–miniGα$_s$ and the atazanavir-stabilized β2AR–βarr1–G$_s$ complexes.**

**(a, b)** Comparison of the PTH1R–βarr1–miniGα$_s$ complex with the atazanavir-stabilized β2AR–βarr1–G$_s$ megaplex (PDB: 9L8L), shown in two orientations to highlight the βarr1–Gα$_s$ interface and the atazanavir-binding site on β2AR.

**(c)** Superposition of the two complexes aligned by their receptors, viewed from two orientations, illustrating the relative positioning of G protein and β-arrestin1.

**(d)** Receptor-aligned superposition comparing the orientation of βarr1 in the two complexes.

**(e, f)** Arrestin-aligned comparison showing the overall β-arrestin 1 conformation (**e**) and detailed views of the finger loop, back loop, and C-loop regions (**f**).

**(g)** Receptor-aligned superposition comparing the position of Gα$_s$ in the two complexes.

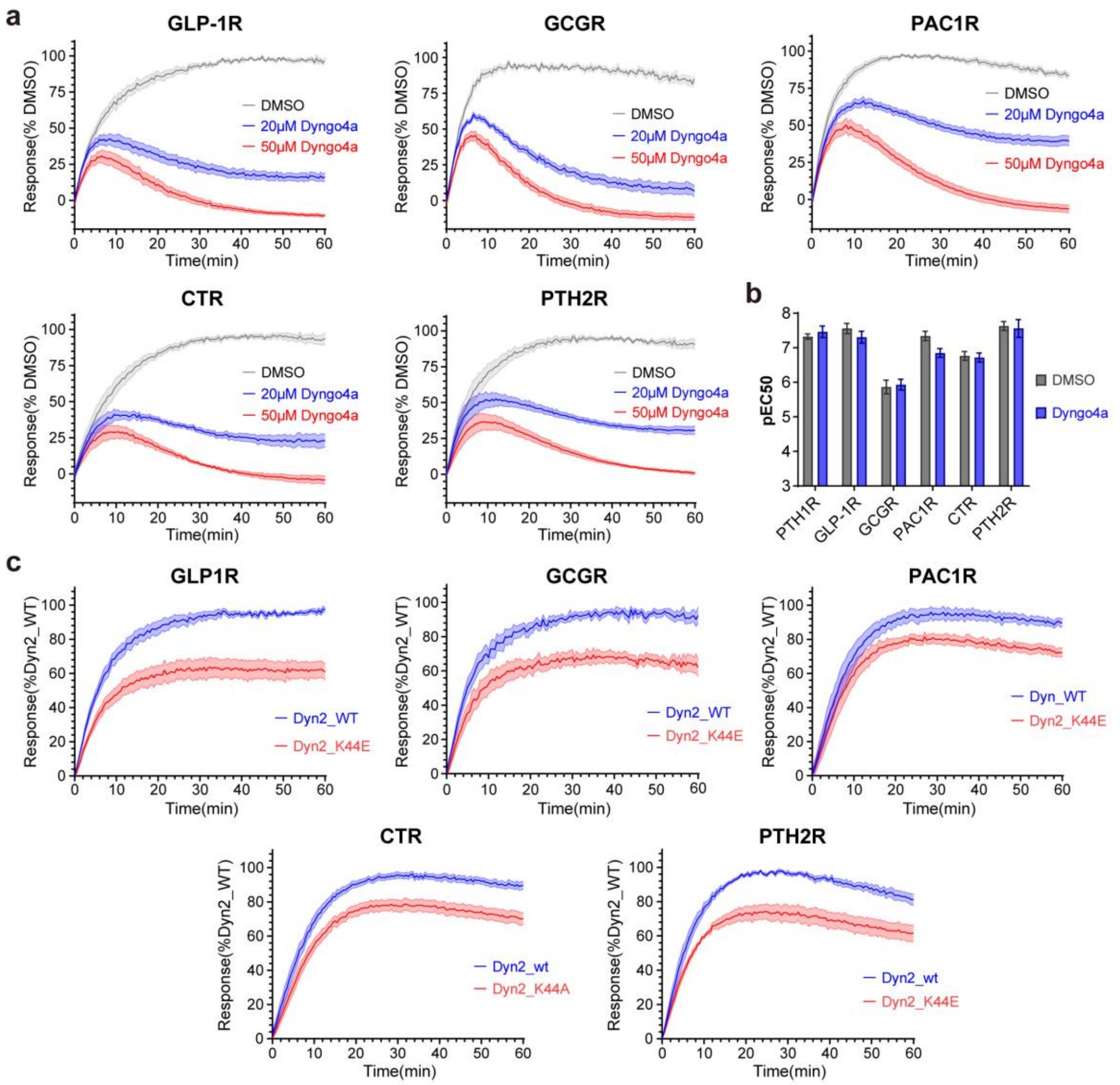


**Extended Data Fig. 8 Inhibition of receptor internalization suppresses sustained cAMP signaling in class B GPCRs.**

(**a**) Dyngo-4a inhibition of cAMP accumulation mediated by class B GPCRs (GLP-1R, GCGR, PAC1R, CTR, PTH2R) upon agonist stimulation, measured by the GloSensor cAMP accumulation assay over a 60-minute period. Luminescence signals were normalized to vehicle-treated control (DMSO).

**(b)** NanoBiT analysis of agonist-induced βarr1 recruitment, with LgBiT fused to receptor C-terminus and SmBiT fused to βarr1 N-terminus. Effects of Dyngo-4a versus vehicle (DMSO) are shown as $pEC_{50}$ in the bar graph.

**(c)** Effect of wild-type or dominant-negative Dynamin2 (Dyn2_K44E) on cAMP accumulation mediated by the same GPCRs, measured by GloSensor cAMP assay over a 60-minute period and normalized to Dyn2_WT. Related to Extended Data Fig.

1c.

Data are presented as mean ± S.E.M. from at least three independent experiments, each performed in triplicate.

**Extended Data Table 1. Functional characterization of different G protein subunits fused with LgBiT.**

| with PTH1R_wt + SmBiT_βarrestin1 | LA-PTH | | Cell surface expression |
|---|---|---|---|
| | $pEC_{50}$±S.E.M. | Span±S.E.M. (%WT) | (% WT) |
| LgBiT_miniG$\alpha_s$+Gβγ | 7.61 ± 0.07 | 100.0 ± 0.0 | 100.0 ± 8.2 |
| miniG$\alpha_s$_LgBiT+Vector | ND | ND | 103.9 ± 2.8 |
| miniG$\alpha_s$_LgBiT+Gβγ | ND | ND | 100.9 ± 11.1 |
| Gβ_LgBiT+Gγ+Vector | ND | ND | 131.7 ± 1.2 |
| Gβ_LgBiT+Gγ+miniG$\alpha_s$ | ND | ND | 103.0 ± 10.2 |
| LgBiT_Gβ+Gγ+Vector | 7.57 ± 0.13 | 18.3 ± 3.1**** | 72.9 ± 2.8 |
| LgBiT_Gβ+Gγ+miniG$\alpha_s$ | 7.77 ± 0.12 | 14.4 ± 1.9**** | 110.3 ± 5.0 |
| Gγ_LgBiT+Gβ+Vector | ND | ND | 103.2 ± 1.2 |
| Gγ_LgBiT+Gβ+miniG$\alpha_s$ | ND | ND | 108.5 ± 10.6 |
| LgBiT_Gγ+Gβ+Vector | ND | ND | 88.1 ± 2.4 |
| LgBiT_Gγ+Gβ+miniG$\alpha_s$ | ND | ND | 84.0 ± 10.1 |

$pEC_{50}$ values, span, and relative expression levels of various G protein subunits fused to LgBiT were assessed. Data were normalized to the maximum and minimum responses of LgBiT–miniG$\alpha_s$ in the presence of Gβγ and presented as mean ± S.E.M. from at least three independent experiments performed in triplicate. Dose–response curves were fitted using a three-parameter logistic model to calculate $pEC_{50}$and span in NanoBiT complementation assays. Expression levels were determined by flow cytometry. Statistical analysis was performed using two-sided one-way ANOVA with Tukey's multiple comparisons test. ND, not detected. *$P < 0.05$; **$P < 0.01$; ***$P < 0.001$; ****$P < 0.0001$.

**Extended Data Table 2. Cryo-EM data collection, model refinement and validation statistics.**

| **Data collection and processing** | |
|---|---|
| Magnification | 165K |
| Voltage (kV) | 300 |
| Electron exposure (e-/Å$^2$) | 50 |
| Defocus range (mm) | -0.8 to -1.8 |
| Pixel size (Å) | 0.73 |
| Symmetry imposed | C1 |
| Final particle images(no.) | 194,863 |
| Map resolution (Å) | 3.1 |
| FSC threshold | 0.143 |
| **Refinement** | |
| Initial model | Alphafold2 |
| Model resolution (Å) | 2.9/4 |
| FSC threshold | 0.143/0.5 |
| Model-Map CC (mask) | 0.63 |
| **Model composition** | |
| Non-hydrogen atoms | 9738 |
| Protein residues | 1208 |
| B factors (Å$^2$) | |
| Protein | 51.78 |
| R.m.s. deviations | |
| Bond lengths (Å) | 0.002 |
| Bond angles (°) | 0.517 |
| **Validation** | |
| MolProbity score | 1.65 |
| Clash score | 5.51 |
| Rotamer outliers (%) | 0.57 |
| Ramachandran plot | |
| Favored (%) | 94.84 |
| Allowed (%) | 4.99 |
| Disallowed (%) | 0.17 |

**Extended Data Table 3. Functional analysis of β-arrestin1 loop substitution mutants.**

| SmBiT_βarrestin1 (with PTH1R_wt + LgBiT_miniG$\alpha_s$) | LA-PTH | | Expression level |
|---|---|---|---|
| | $pEC_{50}$±S.E.M. | Span±S.E.M. (%WT) | (% WT) |
| WT | 7.22 ± 0.09 | 100.0 ± 0.0 | 100.0 ± 3.1 |
| Finger Loop_5G | 7.33 ± 0.11 | 46.9± 2.8**** | 113.1 ± 5.2 |
| C Loop_5G | 7.29 ± 0.10 | 61.7 ± 3.7**** | 118.1 ± 4.8 |
| Back Loop_5G | 7.44 ± 0.11 | 60.1 ± 4.4**** | 131.8 ± 2.7 |

$pEC_{50}$ values, span, and relative expression levels of βarr1 mutants with loop substitutions were evaluated. Data are presented as mean ± S.E.M. from at least three independent experiments performed in triplicate. Dose–response curves were analyzed using a three-parameter logistic equation to calculate $pEC_{50}$ and span through NanoBiT complementation assays. Expression levels were measured by flow cytometry. Statistical significance was assessed by one-way ANOVA with Tukey's multiple comparisons test. *$P < 0.05$; **$P < 0.01$; ***$P < 0.001$; ****$P < 0.0001$.

**Extended Data Table 4. Functional analysis of miniGα$_s$ loop substitution mutants.**

| LgBiT_miniGα$_s$ （with PTH1R_wt + SmBiT_βarrestin1） | LA-PTH | | Expression level |
|---|---|---|---|
| | pEC$_{50}$±S.E.M. | Span±S.E.M. (%WT) | (% WT) |
| WT | 7.12 ± 0.14 | 100.0 ± 0.0 | 100.0 ± 1.7 |
| Switch I_5G | 6.97 ± 0.06 | 66.1 ± 6.6*** | 100.5 ± 7.6 |
| Switch II_5G | 6.98 ± 0.05 | 7.1 ± 2.00**** | 94.8 ± 5.3 |
| α3/β5-loop_5G | 6.93 ± 0.04 | 39.7 ± 5.2**** | 87.6 ± 1.3 |

pEC$_{50}$ values, span, and relative expression levels of miniGα$_s$ mutants with loop substitutions were assessed. Data represent mean ± S.E.M. from at least three independent experiments performed in triplicate. Dose–response curves were fitted using a three-parameter logistic model to determine pEC$_{50}$ and span in NanoBiT complementation assays. Expression levels were quantified by flow cytometry. Statistical analysis was performed using one-way ANOVA with Tukey's multiple comparisons test. *$P < 0.05$; **$P < 0.01$; ***$P < 0.001$; ****$P < 0.0001$.

**Extended Data Table 5. Functional analysis of miniG$\alpha_s$ single-point mutants.**

| LgBiT_miniG$\alpha_s$ (with PTH1R_wt + SmBiT_βarrestin1) | LA-PTH | | Expression level |
|---|---|---|---|
| | pEC$_{50}$±S.E.M. | Span±S.E.M. (%WT) | (% WT) |
| WT | 7.17 ± 0.12 | 100.0 ± 0.0 | 100.0 ± 2.5 |
| K233A | 7.19 ± 0.12 | 64.9 ± 4.7*** | 87.5 ± 3.5 |
| W234A | 7.17 ± 0.13 | 67.4 ± 5.0*** | 101.0 ± 5.7 |
| C237A | 7.25 ± 0.13 | 72.8 ± 2.0** | 105.5 ± 2.6 |
| D240A | 7.29 ± 0.07 | 72.4 ± 6.0** | 100.2 ± 4.2 |
| W281A | 7.32 ± 0.09 | 69.1 ± 5.9** | 95.6 ± 2.1 |

pEC$_{50}$ values, span, and relative expression levels of miniG$\alpha_s$ single-point mutants were assessed. Data represent mean ± S.E.M. from at least three independent experiments performed in triplicate. Dose–response curves were fitted using a three-parameter logistic model to determine pEC$_{50}$ and span in NanoBiT complementation assays. Expression levels were quantified by flow cytometry. Statistical analysis was performed using one-way ANOVA with Tukey's multiple comparisons test. *$P < 0.05$; **$P < 0.01$; ***$P < 0.001$; ****$P < 0.0001$.

**Extended Data Table 6. Functional analysis of βarr1 and miniGα$_s$ loop substitution mutants for GLP-1R.**

| GLP-1R_wt + SmBiT_βarrestin1 + LgBiT_miniGα$_s$ | GLP-1 | | Expression level |
|---|---|---|---|
| | pEC$_{50}$±S.E.M. | Span±S.E.M. (%WT) | (% WT) |
| WT | 7.25 ± 0.10 | 100.00 ± 0.00 | 100.0 ± 3.3 |
| βarr1_Finger Loop_5G | 7.25 ± 0.08 | 58.9 ± 6.0 *** | 103.8 ± 3.7 |
| βarr1_C Loop_5G | 7.26 ± 0.03 | 67.4 ± 7.0** | 111.1 ± 2.3 |
| βarr1_Back Loop_5G | 7.15 ± 0.06 | 74.7 ± 5.9* | 106.4 ± 1.1 |
| miniGα$_s$_Switch I_5G | 7.14 ± 0.08 | 57.4 ± 6.3*** | 109.4 ± 3.9 |
| miniGα$_s$_Switch II_5G | 7.41 ± 0.19 | 3.9 ± 1.0**** | 137.4 ± 2.9 |
| miniGα$_s$_α3/β5 Loop_5G | 7.12 ± 0.08 | 33.7 ± 4.8**** | 88.3 ± 2.0 |

pEC$_{50}$ values, span, and relative expression levels of βarr1 and miniGα$_s$ mutants with loop substitutions were evaluated. Data are presented as mean ± S.E.M. from at least three independent experiments performed in triplicate. Dose–response curves were analyzed using a three-parameter logistic equation to calculate pEC$_{50}$ and span in NanoBiT complementation assays. Expression levels were measured by flow cytometry. Statistical significance was assessed by one-way ANOVA with Tukey's multiple comparisons test. *$P < 0.05$; **$P < 0.01$; ***$P < 0.001$; ****$P < 0.0001$.
